# The ZmHSF20–ZmHSF4–ZmCesA2 module regulates heat stress tolerance in maize

**DOI:** 10.1101/2024.02.21.581499

**Authors:** Ze Li, Zerui Li, Yulong Ji, Chunyu Wang, Shufang Wang, Yiting Shi, Jie Le, Mei Zhang

## Abstract

Temperature shapes the geographical distribution and behavior of plants. Understanding the regulatory mechanisms behind plant heat response is important for developing climate-resilient crops, including maize (*Zea mays*). To identify transcription factors that may contribute to heat response, we generated a dataset of short- and long-term transcriptome changes following a heat treatment time course in the maize inbred line B73. Co-expression network analysis highlighted several transcription factors, including the class B2a heat shock factor ZmHSF20. *ZmHsf20* mutant seedlings exhibited enhanced tolerance of heat stress. Furthermore, DNA affinity purification sequencing and CUT&Tag assays demonstrated that ZmHSF20 binds the promoters of *Cellulose synthase A2* (*ZmCesA2*) and three class A *HSF* genes, including *ZmHSF4*, repressing their transcription. We showed that ZmCesA2 and ZmHSF4 positively regulate heat response, with ZmHSF4 directly activating *ZmCesA2* transcription. In agreement with the transcriptome analysis, ZmHSF20 negatively modulated cellulose accumulation and repressed the expression of cell wall–related genes. Importantly, the *ZmHsf20 ZmHsf4* double mutant exhibited decreased thermotolerance, placing ZmHSF4 downstream of ZmHSF20. Based on our results, we propose an expanded model of the heat stress response in maize, whereby ZmHSF20 lowers heat tolerance of seedlings by repressing *ZmHSF4* and *ZmCesA2*, thus balancing growth and defense at the seedling stage.

**One-sentence summary:** ZmHSF20, as a negative factor, acts upstream of *ZmHSF4* and *ZmCesA2*, which are involved in positively regulating the cell wall development under heat stress, thereby improving maize heat tolerance.

## Introduction

Global warming is responsible for the increasingly frequent occurrence of intense heat waves characterized by very hot days and nights (Sinha et al., 2021). Crops and other plants are greatly impacted by this rise in global average temperatures, which threatens plant growth and thus agricultural yields (Zhao et al., 2017). Maize (*Zea mays*) is one of the three most important food crops worldwide and is severely affected by heat stress (HS), with a 7% value for drop in yield with each 1°C increase in temperature (Tigchelaar et al., 2018). Hence, it is urgent to explore the genes that comprise the regulatory networks behind plant tolerance of HS to produce more climate-resilient varieties of maize, as well as other crops.

Heat shock factors (HSFs) are key regulatory factors of the plant HS response. When plants experience HS, HSFs bind to the cis-regulatory element known as Heat Shock Element (HSE, 5′-nGAAnnTTCn-3′) present in the promoters of downstream genes to regulate the transcription of heat shock protein (*HSP*) genes (Andrasi et al., 2021). HSFs are divided into three subfamilies, A, B, and C, based on their specific structures (von Koskull-Doring et al., 2007), although all HSF members contain a Deoxyribonucleic acid (DNA)-binding domain (DBD) and an oligomerization domain (OD) (Scharf et al., 2012). Among the HSFA subfamily, in Arabidopsis (*Arabidopsis thaliana*), HSFA1 is the master regulator that activates transcriptional networks in response to HS (Yoshida et al., 2011). In addition, SlHSFA7 affects plant heat tolerance by regulating the expression of *SlHSFA1* in tomato (*Solanum lycopersicum*) (Mesihovic et al., 2022). In wheat (*Triticum aestivum*), TaHSFA1 functions in heat tolerance, and interaction with ubiquitin-binding proteins alters TaHSFA1 activity (Wang et al., 2023). Alternative splicing modulates the extent of transcriptional activation in response to HS in wheat, with alternative splicing of *TaHSFA6e* producing two major forms of TaHSFA6e that are distinguished by their binding affinity for downstream *HSP* genes (Wen et al., 2023). The overexpression of *FaHSFA2c* in tall fescue (*Festuca arundinacea*) enhances plant heat tolerance by inducing the transcriptional activation of heat stress responsive genes (Wang et al., 2017).

The HSFC and HSFB subfamilies also have important functions in plants. Arabidopsis plants heterologously expressing *FaHSFC1b* displayed enhanced tolerance of HS compared to control plants, suggesting that FaHSFC1b enhances heat tolerance in Arabidopsis (Zhuang et al., 2018). Moreover, OsHSFC1b contributes to the salt tolerance response in rice (*Oryza sativa*) (Schmidt et al., 2012). Notably, the heat tolerance conferred by HSFB subfamily members varies among plant species. In Arabidopsis, AtHSFB1 and AtHSFB2b are negative regulators of heat tolerance and their expression is essential for acquired heat tolerance (Ikeda et al., 2011). Similarly, Arabidopsis and rice plants heterologously expressing *ZmHSFB2b* displayed decreased tolerance for heat mediated by a mechanism affecting the accumulation of reactive oxygen species (ROS) (Qin et al., 2022). By contrast, in tomato, SlHsfB1 is a coactivator that cooperates with SIHsfA1 to positively regulate plant heat tolerance (Bharti et al., 2004). In grape (*Vitis vinifera*), VvHSFB1 acts as a positive regulator of HS responses (Chen et al., 2023). The regulatory mechanism of HSFs has not, however, been explored in maize.

In addition, there is very little information about the functions of non-HSF proteins responsible for HS in crops, especially in maize as compared to Arabidopsis (Ohama et al., 2017). It is known that HS conditions can reduce protein stability, leading to the accumulation of misfolded proteins in the endoplasmic reticulum (ER) (Howell, 2013). For example, ZmbZIP60 improves the heat tolerance of maize seedlings by increasing the expression of the gene heat up-regulated (*Hug1*), which influences the stability of proteins in the ER (Li et al., 2020; Xie et al., 2022). Overexpression of ZmHsp101 in maize anthers results in robust microspores under HS (Li et al., 2022); similarly, loss of invertase alkaline neutral 6 (ZmInvan6) function affects the progression of maize meiosis at high temperature and lowers pollen fertility, thus influencing yield (Huang et al., 2022). The transcription factor Necrotic upper tips1 (nut1) participates in heat and drought stress responses by directly influencing cellulose biosynthesis and apoptosis during protoxylem development in maize flowering (Dong et al., 2020). The mitogen-activated protein kinase ZmMPK20 is phosphorylated by its upstream kinase MPAPK kinase 9 (ZmMKK9) and regulates the stability of RPM1-interacting protein 2 (ZmRIN2) to enhance maize thermotolerance (Cheng et al., 2023).

Overall, much remains to be investigated about the regulatory mechanism underlying HS response in maize. Here, we identified ZmHSF20 as a central regulator of the maize HS response. We demonstrated that knocking out *ZmHSF20*, overexpressing *ZmHSF4* (a member of the ZmHSFA subfamily), or overexpressing the cellulose synthase gene *ZmCesA2* all improve the heat tolerance of maize seedlings. We defined the two submodules ZmHSF20–ZmHSF4–ZmCesA2 and ZmHSF20–ZmCesA2 as playing central roles in shaping maize heat tolerance mediated by cell wall–related genes. Understanding of these regulatory modules will elucidate the function of *ZmHSF*s in heat stress tolerance in maize.

## Results

### Identification of key transcription factors related to the heat stress response

To identify putative transcription factors (TFs) with a role in the response to heat stress (HS), we performed transcriptome deep sequencing (RNA-seq) of B73 seedlings exposed to heat treatments of 45°C for 5 min, 15 min, 30 min, 2 h, or 8 h, together with matching control seedlings maintained under control conditions. We conducted three replicates for each condition, whose results were all highly correlated (Supplemental Data Set S1). We then performed weighted gene co-expression network analysis (WGCNA) by using differentially expressed genes (DEGs), and classified them into five modules (Figure 1A). Gene Ontology (GO) analysis of these five modules revealed that the turquoise module displays an enrichment in the GO terms ‘response to heat’ and ‘heat acclimation’, which are likely to be directly related to the HS response (Figure 1B and Supplemental Data Set S2). Furthermore, members of the HSF and ERF families are significantly overrepresented in the turquoise module (Figure 1C). We constructed a co-expression regulatory network using the 8 HSF transcription factor genes (five *HSFA* genes and three *HSFB* genes) present in our data, and we observed that ZmHSF20 are the top three HSFs with the largest number of connections (Figure 1D); then, by checking their expression patterns under heat treatment, we found *ZmHSF20* was strongly induced by heat stress (Supplemental Figure S1). Hence, we considered *ZmHSF20* as a potentially essential TF in the HS response. Phylogenetic analysis of the HSF protein families of Arabidopsis, rice, and maize revealed that ZmHSF20 belongs to the HSFB2 subfamily (Supplemental Figure S2). We observed no transcriptional autoactivation activity for ZmHSF20 in yeast (*Saccharomyces cerevisiae*) (Supplemental Figure S3).

**Figure 1.**
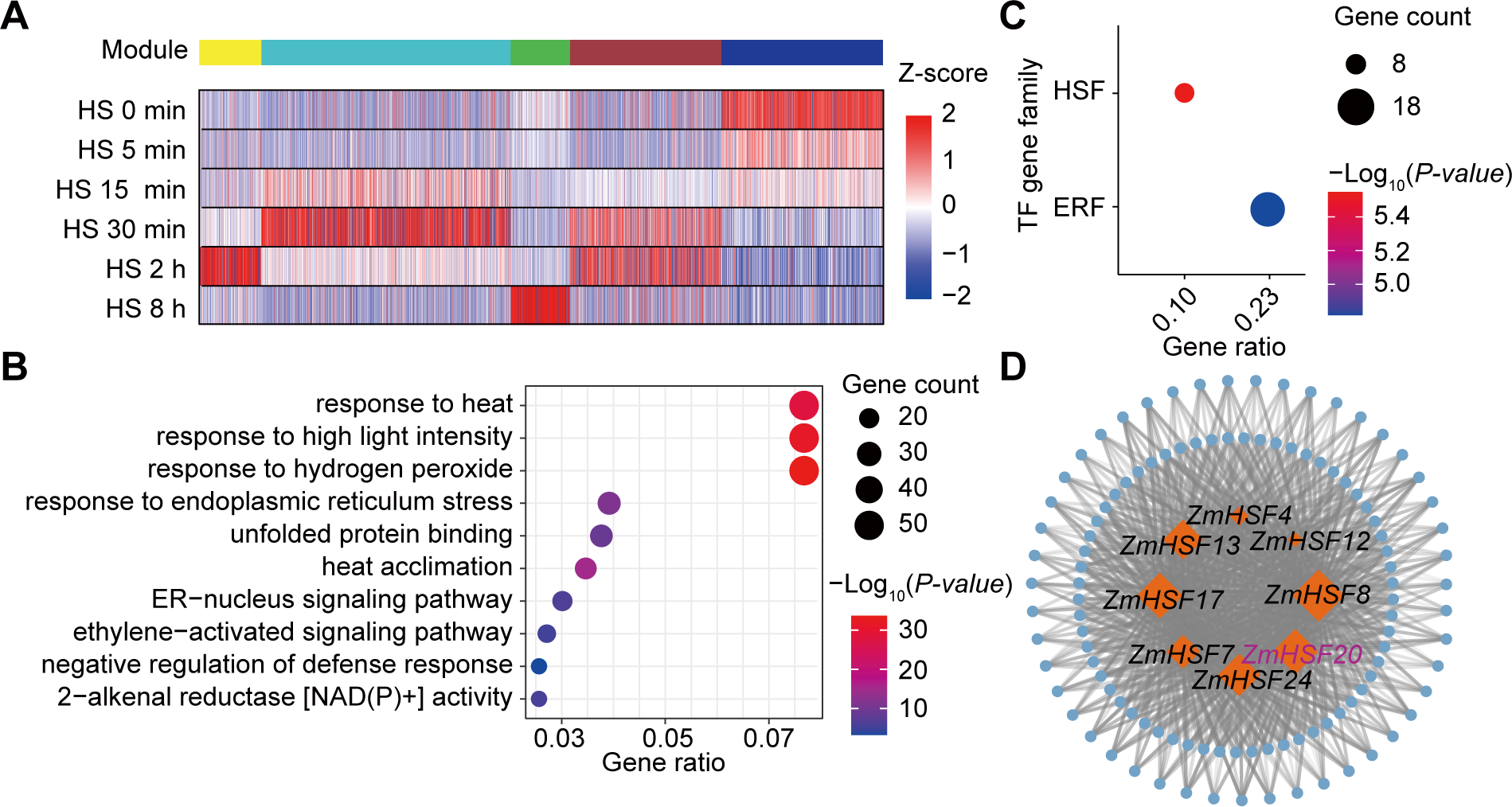
Co-expression analysis and gene regulatory networks of the maize heat stress response constructed by WGCNA. (A) Cluster dendrogram, module detection, and heatmap representation of gene expression of heat response genes at different times during heat treatment. (B) Gene Ontology (GO) term enrichment analysis for genes enriched in the turquoise module. (C) The HSF gene family is significantly overrepresented in the turquoise module. (D) Co-expression network based on eigengenes in the turquoise module. Genes are shown as blue circles, and the gray lines indicate gene–gene associations (false discovery rate < 0.01). The hub genes (transcription factor genes of the *HSF* family) with |weight| > 0.4 are indicated by orange diamonds.

### ZmHSF20 negatively regulates heat tolerance

To explore the function of ZmHSF20 during heat stress, we obtained two knockout lines (*Zmhsf20*-1 and *Zmhsf20-2*) through clustered regularly interspaced short palindromic repeat (CRISPR)/CRISPR-associated nuclease 9 (Cas9)-mediated genome editing. The *Zmhsf20*-1 mutant contains a 252-bp deletion and the *Zmhsf20-2* mutant contains a 24-bp deletion and a 25-bp deletion in the coding region of *ZmHSF20*; both alleles result in a frameshift mutation (Figure 2A). Compared to KN5585 (wild type, WT) controls, both *Zmhsf20* mutants displayed a heat-tolerant phenotype after exposure to 45°C for 3 days followed by a 3-day recovery (Figure 2C), with a survival rate of 50% for the mutants and only about 10% for WT (Figure 2E). We measured higher ROS levels in WT than in the *Zmhsf20* mutants after HS treatment (Figure 2I). Moreover, ion leakage assays indicated that the plasma membrane of *Zmhsf20* mutants suffered less damage upon HS than the WT (Figure 2G). In addition, we assessed the expression level of three classical heat response genes, *ZmHSP20*, *ZmHSP70-1*, and *ZmHSP70-2*. All these genes exhibited significantly increased transcript levels in the *Zmhsf20* mutants relative to the WT (Supplemental Figure S4, A, B and C). To validate the role of *ZmHSF20* in the response to HS, we generated an overexpression line for *ZmHSF20* by driving the *ZmHSF20* coding sequence from the maize *Ubiquitin* (*Ubi*) promoter in the KN5585 background (*ZmHSF20*-OE). We chose two *ZmHSF20*-OE lines with substantially elevated *ZmHSF20* transcript levels, named *ZmHSF20*-OE #1 and *ZmHSF20*-OE #2, for analysis (Figure 2B). Both lines showed a heat-sensitive phenotype compared to WT (Figure 2D), accompanied by increased ion leakage (Figure 2H), and more ROS accumulation (Figure 2 J), and relatively low expression levels of three classical heat response genes (Supplemental Figure S4, D, E and F). Together, these data suggest that ZmHSF20 negatively regulates HS in maize.

**Figure 2.**
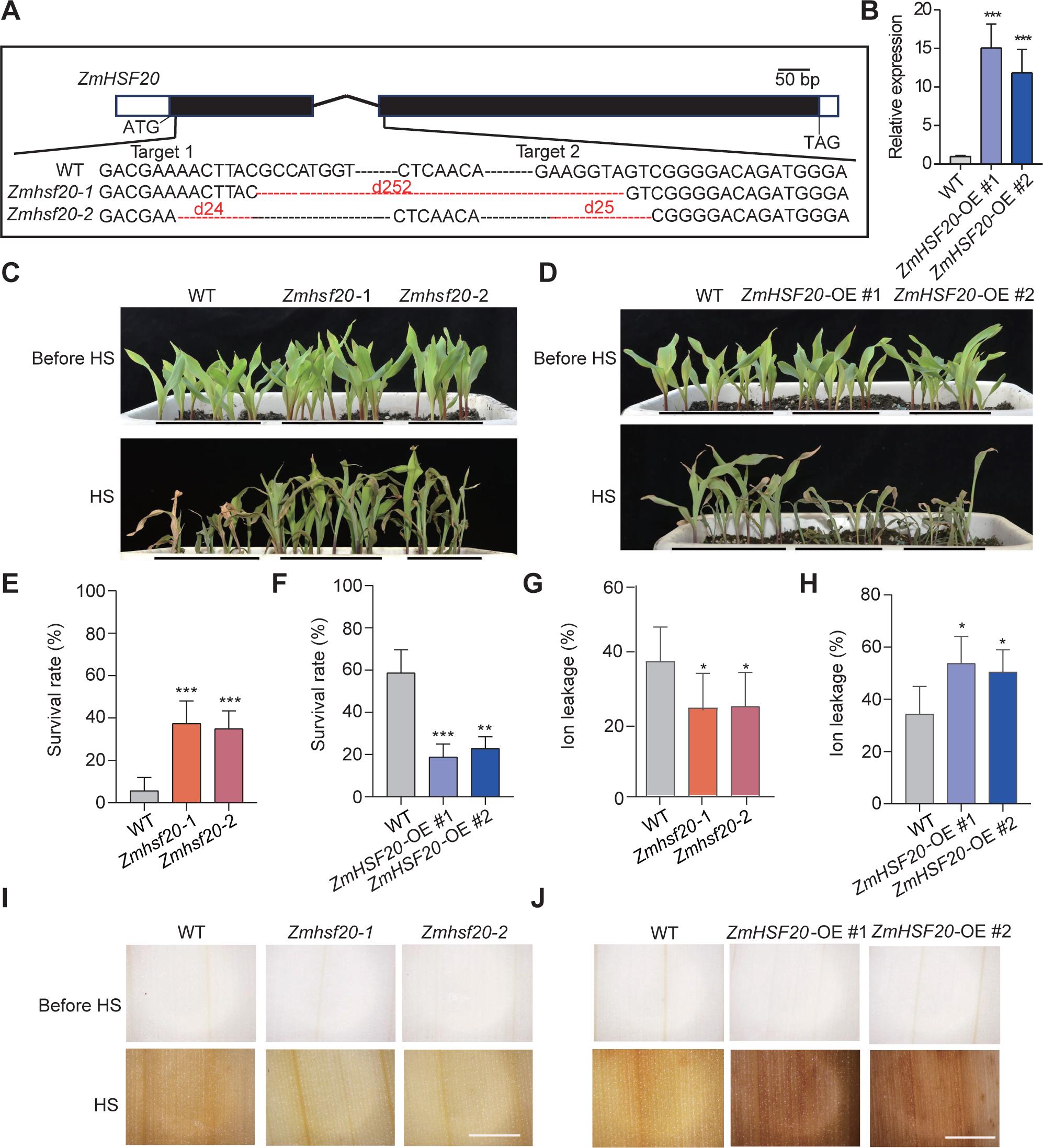
Heat tolerance is modulated by *ZmHSF20*. (A) Construction of CRISPR/Cas9-mediated *Zmhsf20* knockout lines. Two sgRNAs that specifically target *ZmHSF20* were designed, leading to the identification of two mutants, *Zmhsf20*-1 and *Zmhsf20*-2. black rectangles exons, white rectangles Un-Translated Regions (UTRs) and horizontal black lines rectangles introns. (B) Relative *ZmHSF20* transcript levels in the leaves of V2 stage seedlings of *ZmHSF20*-OE and wild type (WT, KN5585). *ACTIN* was used as the internal control. The error bars are based on three independent experiments. The values are means ± SD (n = 3 independent experiments). \*\*\**P* < 0.001, one-way ANOVA. (C) Representative photographs of V2 stage seedlings of WT, *Zmhsf20*-1, and *Zmhsf20*-2 grown at 28L/22L (top) or exposed to 45L for 3 days followed by a 3-day recovery (bottom). (D) Representative photographs of V2 stage seedlings of WT, *ZmHSF20*-OE #1, and *ZmHSF20*-OE #1 grown at 28L/22L (top) or exposed to 45L for 2 days followed by a 3-day recovery (bottom). (E, F), Survival rate of seedlings after recovery at 28L/22L for 3 days in (C, D). The error bars are based on three independent experiments. The values are means ± SD (n = 3 independent experiments). \**P* < 0.05, \*\**P* < 0.01, \*\*\**P* < 0.001, one-way ANOVA. (G, H), Ion leakage rate of V2 stage seedlings grown at 28L/22L exposed to 45L for 1 day. The error bars are based on three independent experiments. The values are means ± SD (n = 3 independent experiments). \**P* < 0.05, \*\*\**P* < 0.001, one-way ANOVA. (I, J) Representative photographs of leaves from V2 stage seedlings grown at 28L/22L, exposed to 45L for 1 day, and stained with DAB, scale bar = 1.5 mm.

To determine the subcellular localization of ZmHSF20, we cloned the full-length coding sequence of ZmHSF20 in frame and upstream of the sequence encoding green fluorescent protein (GFP). We infiltrated the construct into the leaves of *Nicotiana benthamiana* plants and either grew them under normal growth conditions or exposed them to HS treatment. We detected a strong green fluorescent signal in the nucleus in normal and different HS treatments, with full colocalization with the regions stained by a DNA dye, 4’, 6-Diamidino-2-phenylindole dihydrochloride (DAPI) (Supplemental Figure S5). We conclude that ZmHSF20 constitutively localizes in the nucleus regardless of the application of HS.

### ZmHSF20 modulates heat tolerance partly by targeting several *ZmCesA* and ***ZmHSFA* genes**

To identify the direct targets of ZmHSF20, we performed RNA-seq using the *Zmhsf20*-1 mutant and WT. We detected 1,097 differentially expressed genes (DEGs) between the mutant and WT, with 641 genes upregulated and 456 downregulated in *Zmhsf20*-1 mutant (log_2_ fold-change ≥ 1, *P*-value < 0.05, Supplemental Data Set S3). A GO term enrichment analysis revealed that the upregulated genes are primarily enriched in the terms ‘dephosphorylation’, ‘plant-type secondary wall’, ‘cellulose synthase (UDP-forming) activity’, along with several other categories (Figure 3A). In a complementary approach, we conducted DNA affinity purification sequencing (DAP-seq) with recombinant purified ZmHSF20 incubated with an adapter-ligated genomic DNA library in two biological replicates. In total, we detected 77,145 ZmHSF20-binding peaks (*q* < 0.05), of which 4.99% mapped to promoter regions (Supplemental Figure S6). GO analysis indicated that the genes whose promoter regions contain ZmHSF20-binding peaks are primarily enriched in ‘protein folding’, ‘response to heat’, and ‘cellulose synthase (UDP-forming) activity’ (Figure 3B). Indeed, we observed upregulation in the transcript levels of *ZmCesA* genes in the *Zmhsf20* mutants compared to WT, based on reverse-transcription quantitative PCR (RT-qPCR) analysis (Supplemental Figure S7). Taking these results together, we reasoned that *ZmCesA* genes are candidate direct targets of ZmHSF20.

**Figure 3.**
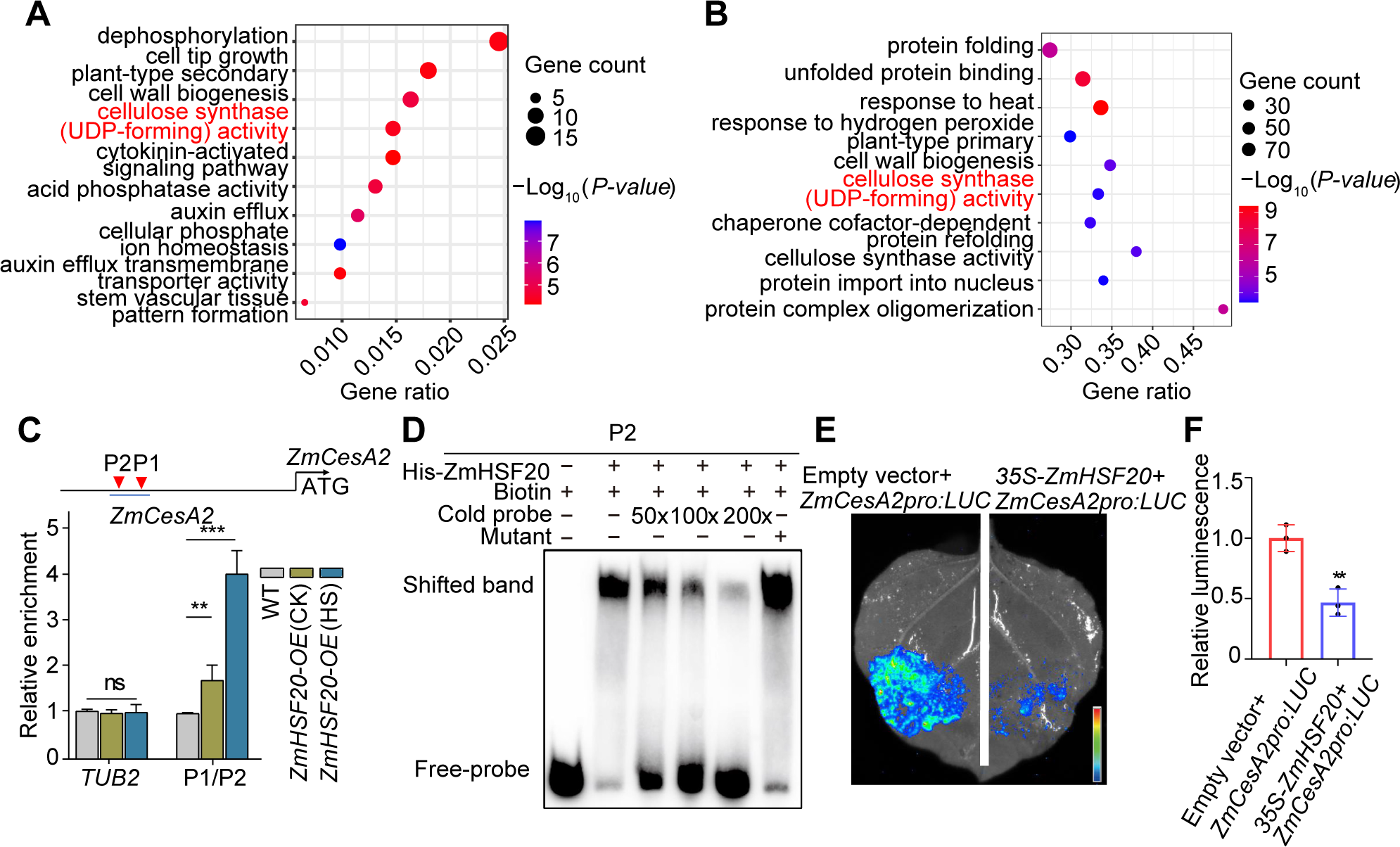
ZmHsf20 binds to the *ZmCesA2* promoter and activates its transcription. (A) GO term enrichment analysis of ZmHSF20-regulated genes following heat stress treatment. (B) Top GO terms enriched among ZmHSF20-bound genes as identified by DAP-seq. (C) CUT&Tag assay showing the binding of ZmHSF20 to the *ZmCesA2* promoter in vivo. WT and *ZmHSF20*-OE seedlings were exposed to heat treatment at 45L for 24 h or maintained at 28L/22L. CUT&Tag was performed using an anti-MYC antibody. Immunoprecipitated DNA was quantified by qPCR using primers specific to regions within the *ZmCesA2* locus. The relative binding of ZmHSF20 to the *ZmCesA2* promoter was normalized to *TUBULIN2* (*TUB2*). Each experiment was performed at least three times with similar results. The values are means ± SD (n = 3 independent experiments). ns, not significant. \*\**P* < 0.01, \*\*\**P* < 0.001, one-way ANOVA. (D) Electrophoretic mobility shift assay (EMSA) showing the binding of recombinant purified ZmHSF20 to the *ZmCesA2* promoter in vitro. The “+” and “−” symbols represent the presence and absence of components, respectively. (E) Interaction assays between ZmHSF20 and the *ZmCesA2* promoter by transient expression in *Nicotiana benthamiana* leaves, based on a luciferase reporter assay. Similar results were obtained for three biological repeats. (F) Quantitative analysis of the LUC activity in (E).

In agreement with this hypothesis, published maize *ZmCesA2* overexpression lines exhibit improved cold tolerance (Zeng et al., 2021). We therefore asked whether ZmHSF20 physically binds to the *ZmCesA2* promoter by performing Cleavage Under Targets and Tagmentation followed by qPCR (CUT&Tag-qPCR) and electrophoretic mobility shift assays (EMSAs). We observed a significant enrichment for the P1/P2 region of the *ZmCesA2* promoter among the immunoprecipitated chromatin by CUT&Tag-qPCR using *ZmHSF20*-OE #1 seedlings and an anti-MYC antibody, especially after HS treatment (Figure 3C). We also used region P2 as a probe for EMSA and detected the specific binding of recombinant ZmHSF20 to a labeled probe containing the P2 region that could be competed away by co-incubation with unlabeled probe (Figure 3D). These results demonstrate that ZmHSF20 binds to the *ZmCesA2* promoter in vitro and in vivo. To validate the effect of ZmHSF20 on *ZmCesA2* transcription, we performed a transient expression assay in *N. benthamiana* leaves by co-infiltrating an effector construct consisting of the full-length *ZmHSF20* coding sequence and the *ZmCesA2* promoter driving the firefly luciferase (*LUC*) reporter gene as reporter construct. We noticed that co-expression of *ZmHSF20* with the *ZmCesA2pro:LUC* reporter decreased LUC activity relative to leaf regions infiltrated only with *ZmCesA2pro:LUC*, indicating that ZmHSF20 directly represses the promoter activity of *ZmCesA2* (Figure 3, E and F). These results indicate that ZmHSF20 suppresses *ZmCesA2* expression by directly binding to its promoter.

Previous studies have shown that some *HSFB* genes affect HS tolerance by modulating the expression of certain *HSFA* genes (Ikeda et al., 2011). However, an interaction between HSFBs and HSFAs in maize has been elusive. To investigate whether ZmHSF20 might regulate *ZmHSFA* family members, we performed a yeast one-hybrid (Y1H) assay to test the binding of ZmHSF20 to the promoters of seven individual *ZmHSFA* genes that are strongly induced by HS treatment (log_2_ fold-change ≥ 2, *P*-value < 0.05, Supplemental Data Set S2, Supplemental Figure S1 B-H). Indeed, we determined that ZmHSF20 binds to the promoters of the *ZmHSFA* genes *ZmHSF4*, *ZmHSF12*, *ZmHSF13*, *ZmHSF14*, *ZmHSF17*, and *ZmHSF24* (Supplemental Figure S8). We used the CUT&Tag-qPCR approach to establish that ZmHSF20 binds only to the promoters of *ZmHSF4*, *ZmHSF12*, and *ZmHSF17* in vivo, with greater binding after heat treatment (Figure 4A and Supplemental Figure S9). These results indicate that ZmHSF20, a member of the HSFB subfamily, may directly bind to the promoters of some *HSFA* subfamily members.

**Figure 4.**
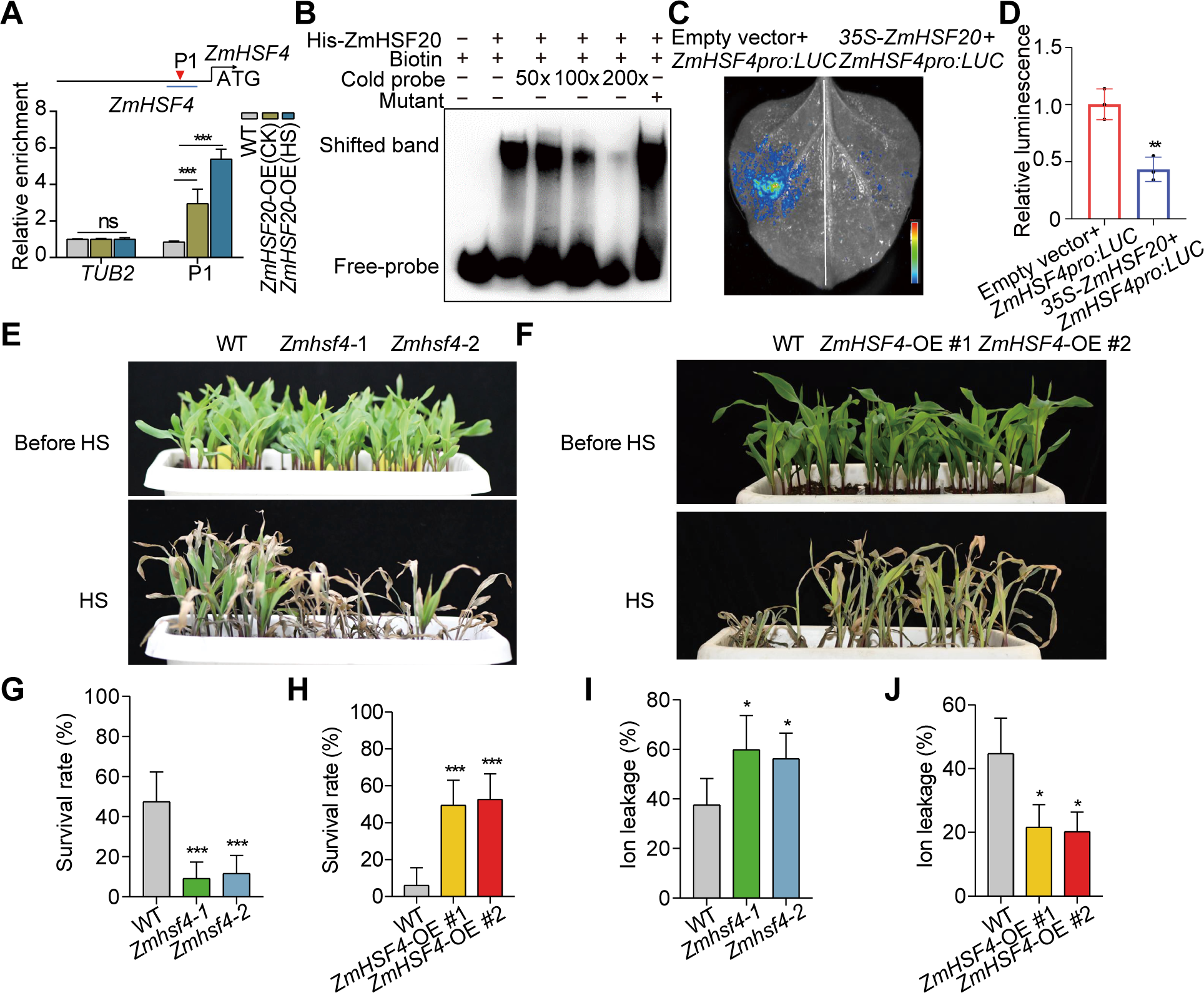
Heat tolerance is modulated by *ZmHSF4*, a downstream target gene of ZmHSF20. (A) CUT&Tag assay showing the binding of ZmHSF20 to the *ZmHSF4* promoter in vivo. WT and *ZmHSF20*-OE seedlings were exposed to heat treatment at 45L for 24 h or maintained at 28L/22L. CUT&Tag was performed using an anti-MYC antibody. Immunoprecipitated DNA was quantified by qPCR using primers specific to regions within the *ZmHSF4* locus. The relative enrichment of ZmHSF20 binding to the *ZmHSF4* promoter was normalized to *TUBULIN2* (*TUB2*). Each experiment was performed at least three times with similar results. The values are means ± SD (n = 3 independent experiments). ns, not significant. \*\*\**P* < 0.001, one-way ANOVA. (B) EMSA showing the binding of recombinant purified ZmHSF20 to the *ZmHSF4* promoter in vitro. The “+” and “−” symbols represent the presence and absence of components, respectively. (C) Interaction assay between ZmHSF20 and the *ZmHSF4* promoter by transient expression in *N. benthamiana* leaves, based on a luciferase reporter assay. Similar results were obtained for three biological repeats. (D) Quantitative analysis of the LUC activity in (C). (E) Representative photographs of V2 stage seedlings of WT, *Zmhsf4*-1, and *Zmhsf4*-2 grown at 28L/22L (top) or exposed to 45L for 2 days followed by a 3-day recovery (bottom). (F) Representative photographs of V2 stage seedlings of WT, *ZmHSF4*-OE #1, and *ZmHSF4*-OE #2 grown at 28L/22L (top) or exposed to 45L for 3 days followed by a 3-day recovery (bottom). (G, H) Survival rate of seedings to recovery at 28L/22L for 3 days in (E, F). The error bars are based on three independent experiments. The values are means ± SD (n = 3 independent experiments). \*\*\**P* < 0.001, one-way ANOVA. (I, J), Ion leakage rate of V2 stage seedlings grown at 28L/22L exposed at 45L for 1 day. The error bars are based on three independent experiments. The values are means ± SD (n = 3 independent experiments). \**P* < 0.05, one-way ANOVA.

### ZmHSF4 positively regulates heat tolerance and functions downstream of ZmHSF20

*ZmHSF4*, like other *ZmHSFA* genes, was highly induced by HS (Supplemental Figure S1H) and belongs to the HSFA2 subfamily (Supplemental Figure S2). We demonstrated that ZmHSF20 binds to the *ZmHSF4* promoter in vitro and in vivo (Figure 4, A and B) and inhibits the transcriptional activity of the *ZmHSF4* promoter in an *N. benthamiana* LUC assay (Figure 4, C and D). From these results, we concluded that ZmHSF20 inhibits *ZmHSFA* transcription.

To explore the role of *ZmHSF4*, we obtained two knockout mutants (*Zmhsf4-1* and *Zmhsf4*-2) via CRISPR/Cas9-mediated genome editing. The *Zmhsf4*-1 mutant harbored a 137-bp deletion and *Zmhsf4*-2 carried a 7-bp deletion in the first exon of *ZmHSF4* (Supplemental Figure S10), both leading to early translation termination. These mutant lines were more susceptible to HS than WT, as evidenced by their lower survival rates (Figure 4, E and G). In agreement, the *Zmhsf4* mutants accumulated more ROS than WT under heat stress but not under normal growth conditions, as determined by DAB staining (Supplemental Figure S11). Similarly, the ion leakage of the *Zmhsf4* mutant lines was much higher than that of WT under heat treatment (Figure 4I). As with ZmHSF20, we determined that ZmHSF4-GFP localizes to the nucleus under both normal and HS conditions (Supplemental Figure S12).

In addition, we generated lines overexpressing *ZmHSF4*. We chose two *ZmHSF4*-OE lines with much higher *ZmHSF4* transcript levels, named *ZmHSF4*-OE #1 and *ZmHSF4*-OE #2 (Supplemental Figure S13). The WT showed a heat-sensitive phenotype compared to *ZmHSF4*-OE #1 and *ZmHSF4*-OE #2 (Figure 4, F and H), which was accompanied by decreased ion leakage and lower ROS accumulation in the overexpression lines (Figure 4J and Supplemental Figure S11). Furthermore, we analyzed the expression level of HS-responsive *HSP* genes in the *Zmhsf4* mutants and *ZmHSF4*-OE plants. The expression levels of *ZmHSP20* and *ZmHSP70* were significantly lower in the *Zmhsf4* mutants and higher in the *ZmHSF4*-OE lines compared to WT following HS treatment (Supplemental Figure S14). These results indicate that ZmHSF4 positively regulates the heat stress response in maize.

To decipher the genetic relationship between *ZmHSF20* and *ZmHSF4*, we generated the *Zmhsf20-1 Zmhsf4-1* double mutant by crossing *Zmhsf20-1* to *Zmhsf4-1*. After 2 days of heat treatment at 45L, the *Zmhsf4-1* and *Zmhsf20 Zmhsf4* double mutants had a survival rate of about 20%, while that of the WT was about 40%. Conversely, the *Zmhsf20-1* mutant displayed strong heat tolerance, with a survival rate at ∼80% (Figure 5, F and G), and the loss of ZmHSF20 function did not suppress the heat-sensitive phenotype of the *Zmhsf4-1* mutant. These genetic results demonstrate that ZmHSF4 functions downstream of ZmHSF20 in the heat stress response.

**Figure 5.**
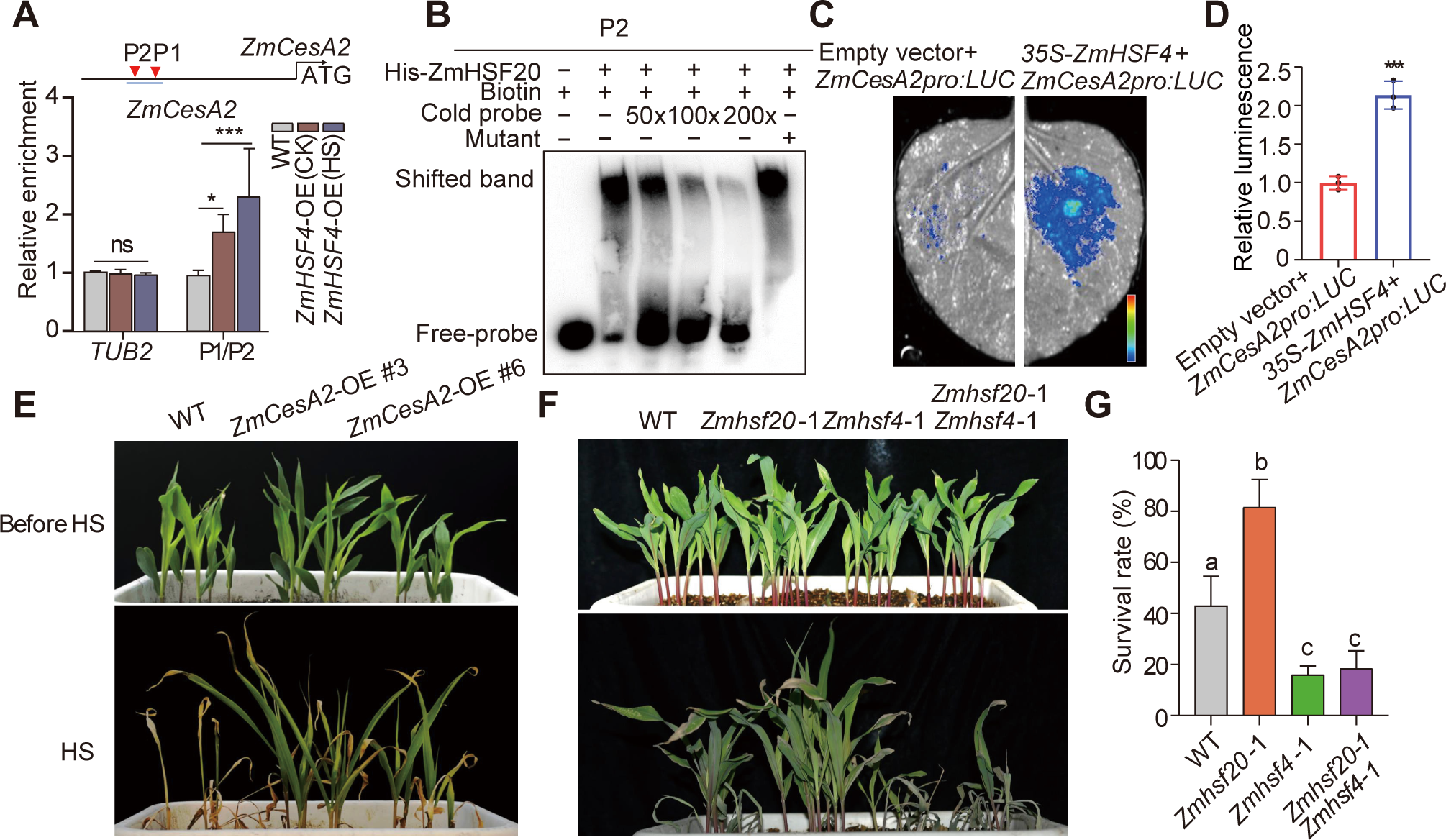
ZmHSF4 binds to the *ZmCesA2* promoter and activates its transcription. (A) CUT&Tag assay showing the binding of ZmHSF4 to the *ZmCesA2* promoter in vivo. WT and *ZmHSF4*-OE seedlings were exposed to heat treatment at 45L for 24 h or were maintained at 28L/22L. CUT&Tag was performed using an anti-MYC antibody. Immunoprecipitated DNA was quantified by qPCR using primers specific to regions within the *ZmCesA2* locus. The relative enrichment of ZmHsf4 binding to the *ZmCesA2* promoter was normalized to *TUBULIN2* (*TUB2*). Each experiment was performed at least three times with similar results. The values are means ± SD (n = 3 independent experiments). ns, not significant. \**P* < 0.05, \*\*\**P* < 0.001, one-way ANOVA. (B) EMSA showing the binding of recombinant purified ZmHsf20 to the *ZmCesA2* promoter in vitro. The “+” and “−” symbols represent the presence and absence of components, respectively. (C) Interaction assay between ZmHSF4 and the *ZmCesA2* promoter in *N. benthamiana* leaves, based on a luciferase reporter assay. Similar results were obtained for three biological repeats. (D) Quantitative analysis of the LUC activity in (C). (E) Representative photographs of WT, *ZmCesA2-*OE #3, and *ZmCesA2-*OE #6 grown at 28L/22L (top) or exposed to 45L for 4 days followed by a 3-day recovery (bottom). (F) Representative photographs of V2 stage seedlings of WT, *Zmhsf20*-1, *Zmhsf4*-1, and *Zmhsf20*-1 *Zmhsf4*-1 grown at 28L/22L (top) or exposed to 45L for 2 days followed by a 3-day recovery (bottom). (G) Survival rate of seedlings after recovery at 28L/22L for 3 days in (H). Different lowercase letters indicate statistically significant differences (adjusted *P* < 0.05, one-way ANOVA).

### *ZmCesA2* is a direct target of ZmHSF4 and confers heat tolerance

As we demonstrated that ZmHSF20 can affects the expression of some *ZmCesA* under HS (Supplemental Figure S7), while *ZmHSF4* is an HSF family member like *ZmHSF20*, we speculated that ZmHSF4 might also affect *ZmCesA* gene expression under HS. We thus analyzed the expression levels of some *ZmCesA* genes in the *Zmhsf4* mutants and WT, which revealed that *ZmCesA2*, *ZmCesA*, and *ZmCesA12* are significantly downregulated in the *Zmhsf4* mutants, compared to WT, upon heat treatment (Supplemental Figure S15). Considering that ZmHSF20 directly targets *ZmCesA2* (Figure 3, C–F), we asked whether ZmHSF4 would also physically interact with the *ZmCesA2* promoter via CUT&Tag-qPCR and EMSA. Indeed, we obtained evidence that ZmHSF4 binds to the *ZmCesA2* promoter in vivo and vitro (Figure 5 A and B). We also co-infiltrated a *LUC* reporter consisting of the *ZmCesA2* promoter driving *LUC* (*ZmCesA2pro:LUC*) together with the empty effector vector (as control) or a *ZmHSF4* effector construct into *N. benthamiana* leaves. We detected low LUC activity from the *ZmCesA2pro:LUC* reporter alone but higher LUC activity when it was co-expressed with *ZmHSF4* (Figure 5, C and D). Overall, these results suggest that ZmHSF4 directly targets *ZmCesA2* and activates its transcription.

To determine whether *ZmCesA* genes regulated by ZmHSF20 and ZmHSF4 contribute to HS tolerance in maize, we obtained two published overexpression lines for *ZmCesA2* (*ZmCesA2-*OE lines), which was previously shown to regulate cold stress in maize (Zeng et al., 2021). Strikingly, *ZmCesA2-*OE lines displayed higher survival rates after HS compared to the WT (LH244) (Figure 5E). These results suggest that ZmCesA2 enhances HS tolerance in maize. Since several studies have shown that there is some correlation between cellulose and heat stress (Le Gall et al., 2015), we measured the cellulose content of WT, the *ZmHSF20*-OE, *ZmHSF4*-OE, and *CesA*-OE lines, and the *Zmhsf20* and *Zmhsf4* mutants. We determined that the *Zmhsf20* mutants and the *ZmHSF4*-OE and *CesA*-OE lines accumulate more cellulose than WT, suggesting that the presence of more cellulose might improve heat tolerance (Supplemental Figure S16).

In a previous study, *ZmCesA* genes and cell wall–related genes were shown to affect the formation of the cell wall, which forms a physical barrier that might influence basic plant tolerance of abiotic stress (Penning et al., 2019). We looked at the transcript levels of several cell wall–related genes mentioned in this study, such as *GALACTURONOSYLTRANSFERASE-LIKE 5* (*ZmGATL5*, Zm00001d028824), a homolog of *Arabidopsis GAUT1* and *GAUT7*, which play essential roles in plant cell wall pectin synthesis (Sterling et al., 2006; Atmodjo et al., 2011); *PHENYLALANINE AMMONIA-LYASE 1* (*ZmPAL1*, Zm00001d017274), a potentially key enzyme involved in the synthesis of cell wall polymer lignin (Huang et al., 2010) and *FERULIC ACID 5-HYDROXYLASE* (*ZmF5H*, Zm00001d032467) which was involved in lignin synthesis (Anderson et al., 2015), in *ZmCesA*-OE, Z*mhsf20*, *Zmhsf4*, and WT (Supplemental Figure S17). All of these genes exhibited increased transcript levels in the *Zmhsf20* mutants, *ZmHSF4*-OE, and *ZmCesA2*-OE. Further, by employing the transmission electron microscope (TEM) analysis, we checked the structure of cell walls in the WT, *Zmhsf20* mutants, and *ZmHSF4-OE* lines. Under HS condition, the structure of cell walls of WT seems to be distorted, while the *Zmhsf20* mutants and *ZmHSF4-OE* lines possessed relatively smooth and uniform cell wall structure; by contrast, no obvious difference was detected among them under normal condition (Supplemental Figure S18). These results together suggest that *Zmhsf20* mutants and *ZmHSF4*-OE lines may directly or indirectly mediate the cell wall development under HS, thereby improving maize heat tolerance.

## Discussion

Only a few *HSF* family genes have been previously reported to be related to abiotic stress in maize, such as *ZmHSF08*, a class A member conferring greater tolerance to salt and drought stress (Wang et al., 2021), and *ZmHSF07* (named as *ZmHsf11*), a class B2b member, whose heterologous overexpression in Arabidopsis and rice led to reduced heat tolerance (Qin et al., 2022). In this study, we report that ZmHSF20, a member of the B2a class of maize HSFs, confers heat tolerance at the seedling stage. Furthermore, ZmHSF20 acts upstream of ZmHSF4 and ZmCesA2, both of which positively regulate heat response by modulating cellulose content and cell wall–related genes expression in maize.

A phylogenetic analysis of HSF proteins from several plant species indicated that ZmHSF20 belongs to the HSFB subfamily (Supplemental Figure S2). Although the HSFB subfamily is the most highly conserved of the three HSF subfamilies, individual members are involved in a variety of biological pathways. In Arabidopsis, the clock protein PSEUDO-RESPONSE REGULATOR 7 (PRR7) regulates rhythmic responses to abiotic stress by repressing the expression of *HSFB2b* (Kolmos et al., 2014). In the Arabidopsis *hsfb1 hsfb2b* double mutant, loss of disease resistance arises through misregulation of the plant defensin genes *PDF1.2a* and *PDF1.2b* (Kumar et al., 2009). In addition, HSFB2b positively regulates salt tolerance in soybean (*Glycine max*) (Bian et al., 2020), reflecting the diversity of HSFB functions and the complexity of their downstream genes. This study adds ZmHSF20 to the small list of ZmHSFB family members that negatively regulate heat tolerance in maize.

In our study, we discovered that the expression of nine of sixteen *ZmHSFA* genes either is not induced by HS or responds only weakly to it, suggesting that these maize HSFAs may be involved in other stress responses and not HS (log_2_ fold-change < 2, *P*-value < 0.05, Supplemental Data Set S2, Supplemental Figure S19). In Arabidopsis, AtHSFA1 is the main factor contributing to heat tolerance (Liu et al., 2011), and AtHSFA2 also is most strongly expressed under HS (Schramm et al., 2006). *AtHSFA2* and *AtHSFA3* largely bind to the same regions in the Arabidopsis genome and may create a transcriptional memory of HS exposure by forming complexes that deposit hypermethylation marks at the promoters of their target genes (Kappel et al., 2023; Friedrich et al., 2021). In addition, Arabidopsis AtHSFB2b and AtHSFB1 are reported to inhibit the expression of *HSFA2* and *HSFA7*, thereby influencing heat tolerance (Kappel et al., 2023). By contrast, our results showed that ZmHSF20, an HSFB2a subfamily member, affects the transcript levels of *ZmHSF12*, *ZmHSF17*, and *ZmHSF4* of the *ZmHSFA2* subfamily (Figure 4A and Supplemental Figure S9). In maize, a previous report indicated that ZmbZIP60, a transcription factor, modulates heat tolerance by regulating *ZmHSF13*, from the *ZmHSFA2* subfamily (Li et al., 2020), while the exact mechanism how ZmHSFA2 regulates heat stress has not been explored. In our study, we demonstrated that ZmHSF4 positively regulate heat tolerance. Therefore, we concluded that the ZmHSFA2 subfamily also has an important function in heat tolerance in maize, albeit ZmHSFA2 targeted by different transcription factors.

The relationship between cellulose synthase and the HS response in maize is unclear. Previous studies revealed that transcription factors from the NAC and MYB families regulate the expression of *CesA* genes (Lampugnani et al., 2019). In Arabidopsis, HSFA7b induces *CesA* expression, thereby affecting salt tolerance (Zang et al., 2019). In this study, we established that members of the ZmHSF family regulate the transcription of *ZmCesA* genes. Overexpression of *ZmCesA2* improved the HS tolerance of maize seedlings. Notably, *ZmCesA2* overexpression also improves cold tolerance in maize (Zeng et al., 2021), suggesting that it may enhance basal tolerance of abiotic stress. Thus, ZmCesA2 and its associated genetic pathway may be important tools for the future engineering of crop plants to resist a range of abiotic stresses.

The plant cell wall is composed of polysaccharides, which protect plants from various external stresses (Wang et al., 2016). Although cell wall remodeling is an important component of plant responses to HS, the exact nature of the connection between cell wall remodeling and HS response is unclear. The structures of pectin and lignin undergo structural changes in response to HS (Wu et al., 2018). Previous studies showed that *CesA* genes are upregulated and cellulose content increases under HS (Le Gall et al., 2015). CesA is crucial in the biosynthesis of cellulose, a major component of plant cell walls. Higher temperatures have been shown to affect the processing speed of CesA, with extremely high temperatures resulting in lower contents of crystalline cellulose (Fujita et al., 2011). In alfalfa (*Medicago sativa*), the expression of several *CesA* genes increases with higher temperature (Guerriero et al., 2014). Thus, *CesA* genes and other cell wall–related genes influence the formation and composition of the cell wall, which in turn would be expected to affect plant stress tolerance (Penning et al., 2019). We speculate that the structure of cell wall change may explain the heat tolerance effects observed in maize. In our study, *ZmCesA* genes were upregulated in *Zmhsf20* mutants and *ZmHSF4*-OE lines under HS conditions (Supplemental Figure S7 and Supplemental Figure S15) leading to increased cellulose production (Supplemental Figure S16). Moreover, the expression of some cell wall–related genes were upregulated between *Zmhsf20* mutants, *ZmHSF4*-OE lines and WT (Supplemental Figure S17). Correspondingly, the structure of cell walls of *Zmhsf20* mutants and *ZmHSF4*-OE lines seems to be more robust to adapt to heat stress compared to WT (Supplemental Figure S18). Nevertheless, the exact mechanism how ZmCesA2 regulates heat stress is to be investigated.

The *Zmhsf20* mutants displayed enhanced tolerance of heat stress compared to WT at the seedling stage, but seed setting rate was comparable between these genotypes at the mature stage. We hypothesize that ZmHSF20 may promote maize growth by repressing heat tolerance during the seedling stage. We did not observe any clear differences in the seed-setting rates of the mutants and WT in plants grown at Yazhou, Sanya, Hainan, which experienced normal temperatures during the winter of 2022, or at Zhuozhou, Hebei, where plants were exposed to HS in July 2023 (http://data.cma.cn/) (Supplemental Figure S20 and Supplemental Figure S21). However, a 1-year experiment is not sufficient to draw strong conclusions. Hence, these experiments should be repeated in the future.

We conclude that ZmHSF20, a member of the maize HSFB subfamily, may decrease heat tolerance by negatively regulating the expression of cellulose synthase–related genes (Figure 6). HSFs regulate plant cell wall remodeling, thereby providing targets and potential strategies for breeding to facilitate plant adaptation to heat stress.

**Figure 6.**
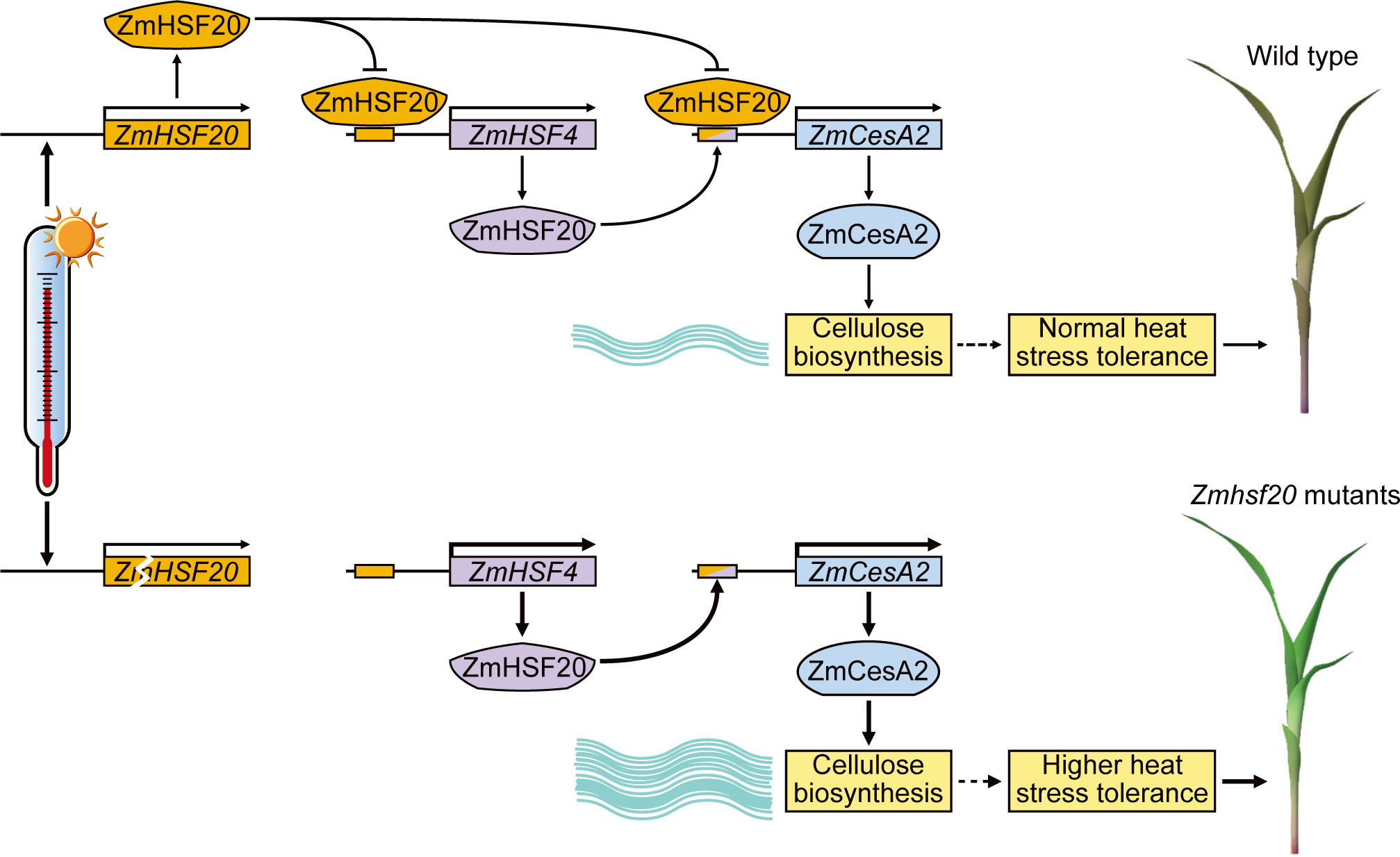
Proposed model of ZmHSF20 function in conferring tolerance to heat stress in maize. Under heat stress, *ZmHSF20* transcripts and ZmHSF20 protein accumulate, leading to the direct transcriptional repression of *ZmHSF4* and *ZmCesA2* expression. In parallel, ZmHSF4 normally promotes the expression of *ZmCesA*s, such that heat treatment further decreases cellulose content in a ZmHSF20- and ZmHSF4-dependent manner.

## Materials and methods

### Plant growth conditions

All transgenic maize (*Z*ea *mays*) plants and CRISPR/Cas9 mutants were generated by WIMI Biotechnology Co. Ltd. The constructs were transformed into the inbred line KN5585. Maize seeds were planted in pots (40 cm × 30 cm × 15 cm, length × width × depth) containing vermiculite, and Pindstrup soil mix (1:1, v:v) and grown at 28L/22L (day/night) under a 16-h-light/8-h-dark photoperiod with 150 µmol m^−2^ s^−1^ white light and 40% relative humidity. For heat treatment, T_2_ generation maize seedlings were grown in pots (40 cm × 20 cm × 15 cm, length × width × depth) to the V2 stage with irrigation before being exposed to 45L under a 16-h-light/8-h-dark photoperiod for 2–4 days using a heat chamber. The control plants were grown in the similar chamber except for heat treatment. After heat treatment, the seedlings were allowed to recover at 28L/22L (16-h-light/8-h-dark) for 3 days prior to imaging.

### Physiological assays

The ion leakage assays were performed as described previously (Zeng et al., 2021). Leaves from V2 stage maize seedlings then being exposed to 45L 1 day under a 16-h-light/8-h-dark) were placed in 15-mL centrifuge tubes with 10 mL double distilled water (ddH_2_O). The solution was vacuumed for 30 min, and the conductance of the water was measured as S0. After being shaken at room temperature for 1 h, the solution was detected as S1. Then the samples were boiled for 30 min and shaken at room temperature for cooling, followed by detection of the conductance of the water, defined as S2. The results were calculated as follows: ion leakage (%) = (S1 − S0)/(S2 − S0).

To detect H_2_O_2_ accumulation in situ, staining with 3’3’-diaminobenzidine (DAB) was used as described by Jambunathan (Jambunathan, 2010). Briefly, V2 stage maize seedlings were exposed to 45L for 1 day before being transferred to 1.0 mg/mL DAB solution for 10 h and vacuum-infiltrated at 37L for 30 min. Subsequently, the leaves were cleared with 95% ethanol until colorless.

### Plasmid construction and plant transformation

The transgenic plants (*ZmHsf20*-OE #1, *OZmHsf20*-OE #2, *ZmHsf4*-OE #1, and *ZmHsf4*-OE #2) used in this study were derived from maize inbred line KN5585. These constructs were generated by cloning the full-length *ZmHsf20* and *ZmHsf4* coding sequences individually into the pBCXUN vector (Zeng et al., 2021). To obtain mutants in *ZmHsf20* and *ZmHsf4* using CRISPR/Cas9-mediated genome editing, fragments of the intron were selected as target for single guide RNA (sgRNA), which were cloned into pBCXUN (Xing et al., 2014). Transgenic plants were obtained by Agrobacterium (*Agrobacterium tumefaciens*) mediated transformation as previously described (Frame et al. 2011).

The full-length coding sequence of *ZmHsf20* was cloned into the pGADT7 vector to generate the AD-*ZmHsf20* construct. The *ZmHsf4*, *ZmHsf11*, *ZmHsf12*, *ZmHsf13*, *ZmHsf14*, *ZmHsf17*, and *ZmHsf24* promoters were individually amplified from KN5585 genomic DNA and cloned into the pHIS2 vector to generate the recombinant pHis2-promoter construct. The pET28a vectors were used to produce His-ZmHsf20 and His-ZmHsf4 recombinant proteins. To obtain the Super:*ZmHsf20*-*GFP* and *ZmHsf4*-*GFP* plasmids, the full-length coding sequences of *ZmHsf20* and *ZmHsf4* were individually cloned into the pSuper:1300-GFP vector. The 2.0-kb promoter fragments of *ZmHsf4*, *ZmCesA*, and *ZmCesA2* were amplified and cloned into the pGreenII 0800-LUC vector to obtain the *ZmHsf4pro:LUC*, *ZmCesApro:LUC*, and *ZmCesA2pro:LUC* constructs, respectively. The *ZmHsf4pro:LUC*, *ZmCesApro:LUC*, and *ZmCesA2pro:LUC* plasmids were transformed separately into Agrobacterium (strain GV3101) together with the helper plasmid (pSoup-P19) for subsequent assays. All constructs were generated using a Seamless Assembly Cloning kit (Clone Smarter). The primers and restriction enzymes used for plasmid construction are listed in Supplemental Data Set S4.

### RNA extraction and RT-qPCR

Total RNA was extracted from the leaves of V2 stage maize seedlings using a plant RNA kit (ER301; TransGen Biotech, China). Total RNA (1µg) was reverse transcribed into first-strand complementary DNA (cDNA) using M-MLV reverse transcriptase according to the manufacturer’s instructions. qPCR analyses were performed as previously described (Zhang et al., 2019). *ACTIN* was used as internal reference to normalize the expression value of each sample. Primers used for qPCR are shown in Supplemental Data Set S4. The experiments were independently performed at least three times (three biological replicates from different plants).

### RNA-seq analysis

For RNA-seq, the leaves and stems of V2 stage maize B73 seedlings grown at 28L/22L (day/night) or exposed to heat treatment at 45L for 5 min, 15 min, 30 min, 2 h, or 8 h, with matching control samples grown at 28L/22L (day/night) for each heat treatment time point were collected to extract total RNA with TRIzol reagent. For RNA-seq, the second leaves of V2 stage maize seedlings (*Zmhsf20*-1 and WT) exposed to heat at 45 L for 24 h were used to extract total RNA with TRIzol reagent. The sequencing library was constructed using 1 µg total RNA from each sample. Libraries were prepared using the TruSeq RNA Library Preparation kit (Illumina, USA) and sequenced on an Illumina Novaseq platform, and 150-bp paired-end reads were generated. Fastp (Chen et al., 2018) was used to remove the adaptor, low-quality bases (Q30), and sequences containing >10% undetermined bases. The filtered reads were aligned to the maize reference genome B73_AGPv4 (Jiao et al., 2017) using Hisat2 (Kim et al., 2015) with default parameters. Subsequently, Stringtie (Pertea et al., 2015) was used to quantify the unique alignment reads (FPKM), and the Pearson correlation coefficient was used to evaluate the repeatability between samples.

### DAP-seq analysis

DAP-seq was performed by Bluescape Hebei Biotech using genomic DNA (gDNA) purified from the leaves of 30-day-old B73 seedlings with a FastPure Gel DNA Extraction Mini Kit (Vazyme) and sonicated to ∼ 200-bp fragments. After end-repair and adenylation, the fragmented gDNA was constructed into sequencing libraries using a NEXTFLEX Rapid DNA Seq Kit (PerkinElmer, Inc., Austin, TX, USA). The full-length coding sequence of *ZmHsf20* was cloned into the pFN19K HaloTag T7 SP6 Flexi vector, and the recombinant protein was produced using a TNT SP6 Coupled Wheat Germ Extract System (Promega, Madison, WI, USA). Magne Halo Tag Beads (Promega) were used to purify and capture the produced protein. The ZmHsf20-bound beads were incubated with the adapter-ligated gDNA library. The beads were then washed 3 times,and the eluted DNA was ligated to an indexed adaptor for sequencing analysis. The eluted DNA from the beads was sequenced on an Illumina NavoSeq6000 instrument with two biological duplicates. The negative input control consisted of Magne Halo Tag Beads incubated with adapter-ligated gDNA library. DAP-seq reads were aligned to the reference genome (B73_AGPv4). For peak analysis, the mapped reads and peak files were examined an using integrative genomics viewer. Enriched motifs were identified by MEME motif discovery software (http://meme-suite.org/). Target genes were defined as those that contained DAP-seq peaks located within 2 kb upstream of the ATG.

### Phylogenetic analysis

Amino acid sequences of HSF family proteins from the plant genome database website (https://phytozome.jgi.doe.gov/pz/portal.html, Supplemental File 1), identified in maize, rice, and Arabidopsis, were aligned using MEGA X software with default pairwise and multiple alignment parameters (Supplemental File 2). The phylogenetic tree was reconstructed based on the alignment results using the neighbor-joining method in MEGA X software with the following parameters: Poisson correction, complete deletion, uniform rates, and bootstrap.

### Protein subcellular localization

The *ZmHsf20-GFP* and *ZmHsf4*-*GFP* constructs were individually introduced into Agrobacterium strain GV3101. The bacterial cultures were resuspended in infiltration buffer (0 mM MgCl_2_, 10 mM MES, pH 5.6, and 150 µM Acetylsyringone) at an OD at 600 nm of 0.8, and then infiltrated into the leaves of *N*. *benthamiana* plants via Agrobacterium (strain GV3101)-mediated infiltration. The infiltrated plants were cultured at 22°C for 3 days before being exposed to heat stress. Confocal imaging and colocalization analysis of the infiltrated leaves were performed using a Zeiss 980 confocal laser scanning microscope with an excitation wavelength of 488 nm and emission wavelengths of 505–530 nm for green fluorescent protein (GFP) and an excitation wavelength of 405 nm and emission wavelengths of 350–450 nm for DAPI.

### Yeast one-hybrid assay (Y1H)

The Y1H assays were conducted according to the manufacturer’s instructions (Matchmaker Gold Y1H Library Screening System; Clontech Laboratories, Mountain View, CA, USA). The pGADT7 empty vector with the pHis2 promoter and pGADT7-ZmHsf20 with the pHis2 empty vector were used as negative controls. The plasmids were cotransformed Y187 Gold yeast strain and plated on synthetic defined (SD) medium without tryptophan, leucine, and histidine (SD−Trp/−Leu/−His). Transformants were plated onto SD−Trp/−Leu/−His medium alone or containing 50 mM 3-amino-1,2,4-triazole (3-AT) as serial dilutions in ddH_2_O.

### Transactivation activity assay

For transactivation assays, the full-length coding sequence of *ZmHsf20* was inserted into the pGBKT7 vector. The resulting plasmid and the empty (pGBKT7) vector were individually transformed into yeast strain AH109 (coolaber) according to the manufacturer’s instructions. Then, positive transformants were selected on SD/−Trp medium and cultured at 30°C for 3 days. Transformants were then serially diluted and spotted onto SD/−Trp/−His/−Ade plates; transactivation activity was assessed based on cell growth.

### Electrophoretic mobility shift assay (EMSA)

Recombinant His-ZmHsf20 and His-ZmHsf4 proteins were produced in *Escherichia coli* strain BL21 (TransGen Biotech). The recombinant proteins were purified with His-tag Purification Resin (TransGen Biotech) according to the manufacturers’ instructions. The purified fusion protein and biotin-labeled DNA probes containing HSE cis-elements were incubated in EMSA/Gel-Shift buffer at at 4L for 1h. EMSAs were carried out using a Light Shift Chemiluminescent EMSA Kit (GS009; Shanghai Beyotime Biotech Co. Ltd) according to the manufacturer’s instructions.

### CUT&Tag-qPCR

The second leaves of *ZmHSF20*-OE and *ZmHSF4*-OE V2 stage seedlings were collected for DNA library construction following the instructions for the Hyperactive Universal CUT&Tag Assay Kit for Illumina (TD903; Vazyme). The leaves collected following the instructions for the Hyperactive Universal CUT&Tag Assay Kit. An anti-MYC antibody was used to immunoprecipitate the protein-DNA complex. The enrichment of DNA fragments was determined by qPCR. All oligonucleotide sequences used are listed in Supplemental Data Set S4.

### Luciferase complementation imaging assays

The full-length coding sequences of *ZmHsf20* and *ZmHsf4* were individually cloned into the pGreenII 62-SK vector as effector constructs. The 2-kb promoter sequences of *ZmHsf4*, *ZmCesA*, and *ZmCesA2* were cloned into the pGreenII-0080-LUC vector to generate the reporter constructs. All effector and reporter constructs were separately transformed into Agrobacterium strain GV3101 (already transformed with pSoup). Then, the appropriate pairs of Agrobacterium cultures harboring an effector plasmid or a reporter plasmid resuspended in infiltration buffer (0 mM MgCl_2_, 10 mM MES, pH 5.6, and 150 µM acetylsyringone) to an OD at 600 nm of 0.8 were co-infiltrated into *N*. *benthamiana* leaves. A low-light, cooled CCD imaging apparatus was used to capture the LUC signal. Relative luminescence was quantified by measuring the intensity with Image J software (Bian et al., 2020). At least six independent leaves were used for each experiment, and three biological replicates from different plants were performed with similar results.

### Cellulose content analysis

Cellulose content assay was carried out according to the protocols described in the Solarbio kit (BC4285). To measure the cellulose contents, the sample is dried at 80L to constant weight, crushed, weighing about 300 mg. Next, extraction solution 1 is added, the samples are placed in a 90 L water bath with 20 min and then centrifuged 6000g at room temperature for 10 min, and the supernatant is discarded. The precipitate is washed twice with 1.5 mL extraction solution; the resulting cleaned precipitate represents the coarse cell wall fraction. The crude cell wall fraction is soaked in 1 mL of extraction solution 2 for 15 hours and then centrifuged 6000g at room temperature for 10 min; the supernatant is discarded, and the precipitate is washed twice with distilled 25L 0.5 mL water and then air dried to obtain the cell wall substance. This is thoroughly homogenized in 0.5mL distilled water by vortexing. Next, 0.75 mL of concentrated sulfuric acid is slowly added and mixed in well, the supernatant is removed and diluted 20 times with distilled water, and the absorption value at 620 nm is determined.

### Transmission electron microscopy

The ultrastructure of maize leaf cell walls in vascular bundles was detected by transmission electron microscopy. The leaf tissues were cut into 1.0 * 3.0 mm pieces and then fixed in 3% (w/v) glutaraldehyde in 0.1 M phosphate buffer solution (PBS) (pH 7.2) at 4L overnight. The fixed tissues were washed in PBS for three times for 20 min each at 4L, post-fixed for 4 h in 1% osmium tetroxide, dehydrated in a graded series of acetone, then infiltrated with Spurr resin (SPI, SPI Chem, West Chester, USA), and polymerized at 60L for 24 h. The samples were cut into ultrathin sections (80-nm thickness-), and examined with a Hitachi transmission electron microscope (H-7700; Hitachi, Japan) at 80 kV.

### Statistical methods

Differences between two groups were assessed using a two-sided Student’s t test. For multiple comparisons, significance analysis was performed with one-way ANOVA followed by Tukey’s multiple comparison tests. The statistical analysis was performed using GraphPad Prism version 8.0 (Supplemental Data Set S5).

## Data availability

The data generated in this study has been uploaded to the NCBI database and can be retrieved under accession number PRJNA1046592. All scripts used for quantification, normalization and statistical testing, are available in the Github: https://github.com/lizerui99/ZmHSF20-ZmHSF4-ZmCesA2.git.

## Accession numbers

Genes mentioned in the study are as follows: *ZmHSF20* (Zm00001d026094), *ZmHSF4* (Zm00001d018941), *ZmHSF12* (Zm00001d046204), *ZmHSF17* (Zm00001d033987), *ZmHSF11* (Zm00001d034433), *ZmHSF24* (Zm00001d032923), *ZmHSF13* (Zm00001d027757), *ZmHSF14* (Zm00001d028269), *ZmCesA2* (Zm00001d037636), *ZmHSP20* (Zm00001d003554), *ZmHSP70-1* (Zm00001d013507), *ZmHSP70-2* (Zm00001d037717), *ZmGATL5* (Zm00001d028824), *ZmPAL1* (Zm00001d017274), *ZmACTIN* (Zm00001d010159) and *ZmF5H* (Zm00001d032467).

## Acknowledgements

We gratefully thank Dr. Feng-Qin Dong (Institute of Botany) and Dr. Jing-Quan Li (Institute of Botany) for technical assistance. This work was supported by the National Key Research and Development Project (2020YFA0509901 and 2020YFE0202300 to M.Z.)

## Author contributions

M.Z. conceived the project and designed the experiments. Z.L. carried out most of the experiments; Z.R.L. performed most data analyses; L.J. collected materials and carried out some experiments. C.W. carried out CUT&Tag. Z.L. and Z.R.L. wrote the draft. J.L., S.W., and Y.S., gave invaluable input to the manuscript. M.Z revise the manuscript. All authors have read and approved the final manuscript.

