## Supplementary material for "The ZmHSF20–ZmHSF4–ZmCesA2 module regulates heat stress tolerance in maize": Supplemnet fIGURE 1-21

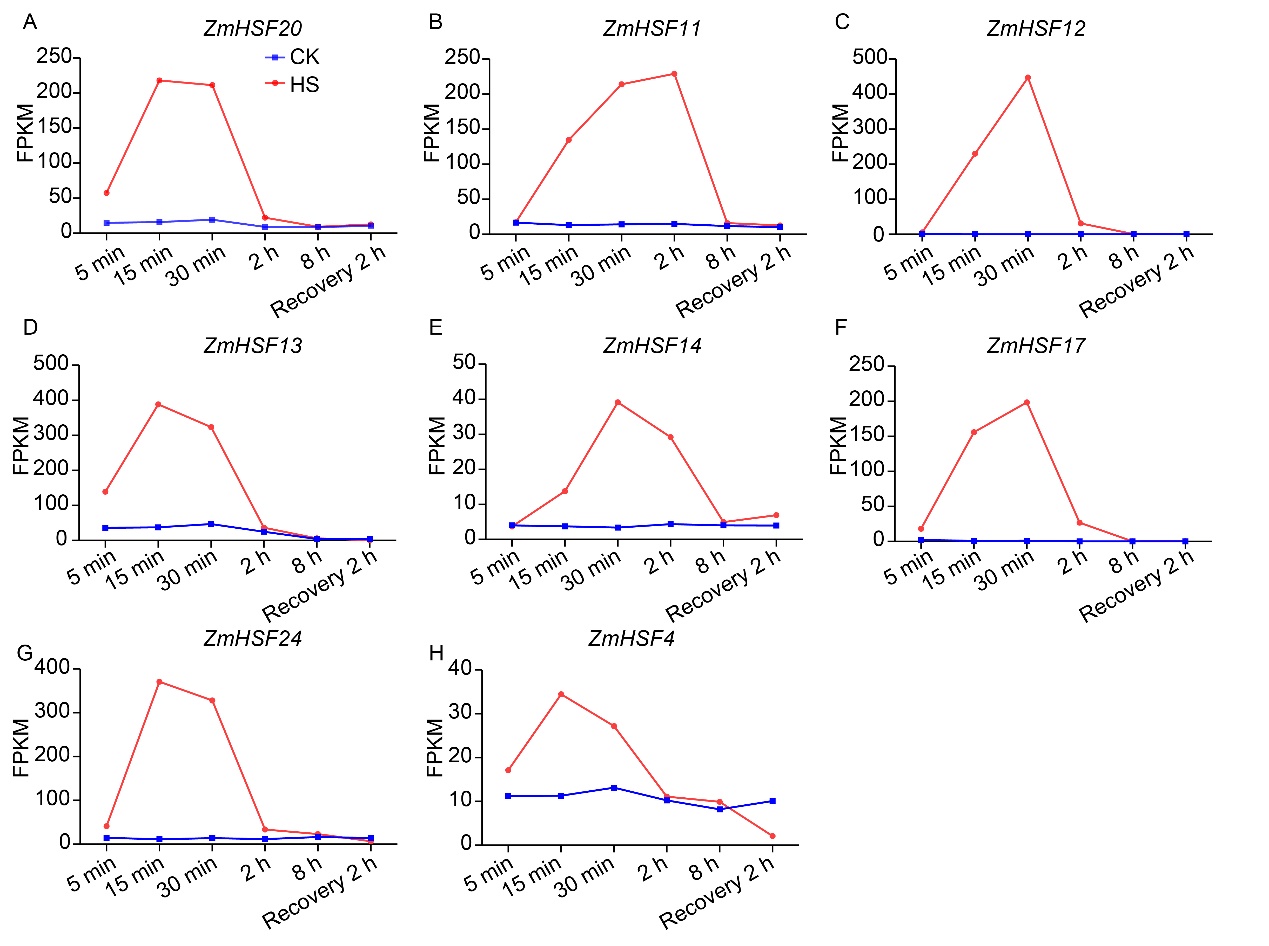

**Supplemental Figure. S1.** *ZmHSF* genes expression under 45℃ treatment response strongly. (Supports Figure 1 and Figure 5) (A-H) Expression pattern of *ZmHSF20* *ZmHSF11,* *ZmHSF12*, *ZmHSF13*, *ZmHSF14*, *ZmHSFf17*, *ZmHSF24*, and *ZmHSF4* during heat treatment.

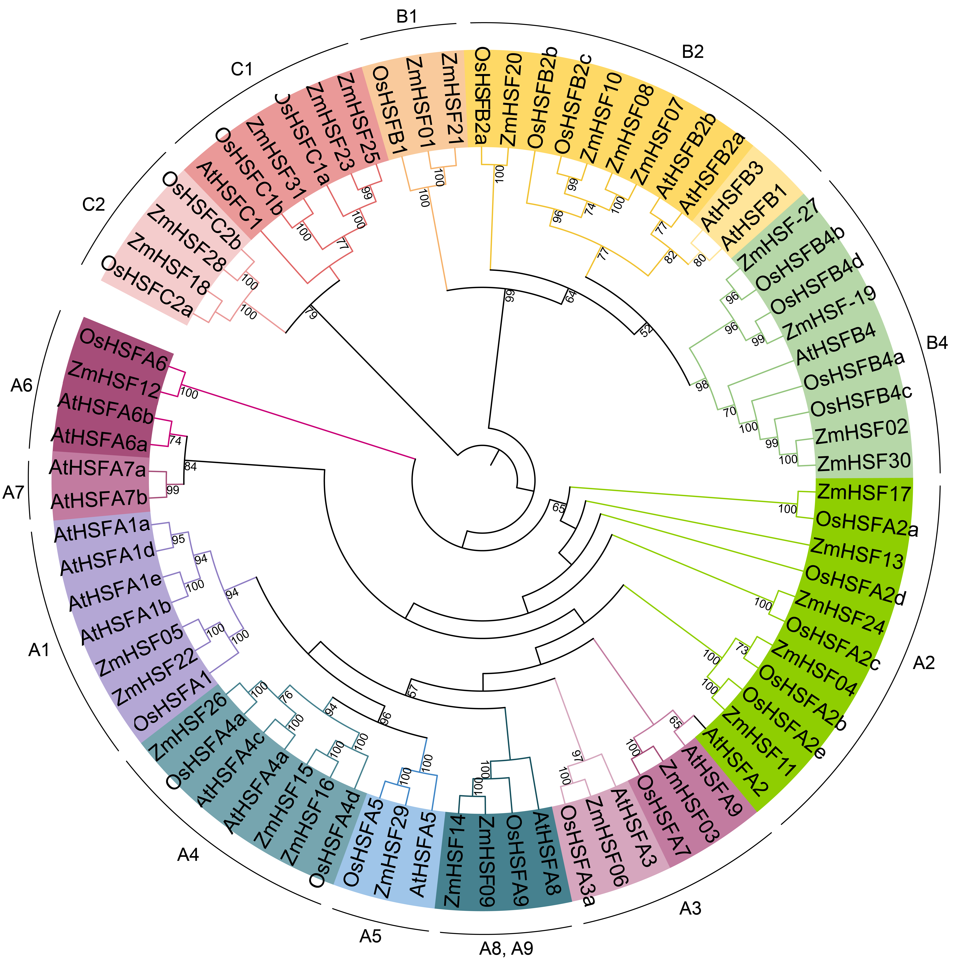

**Supplemental Figure. S2.** Sequence and phylogenetic analyses of HSF family. (Supports Figure 1) Phylogenetic tree reconstructed based on the amino acid sequences of the HSF family members of Arabidopsis, rice, and maize using MEGA X software.

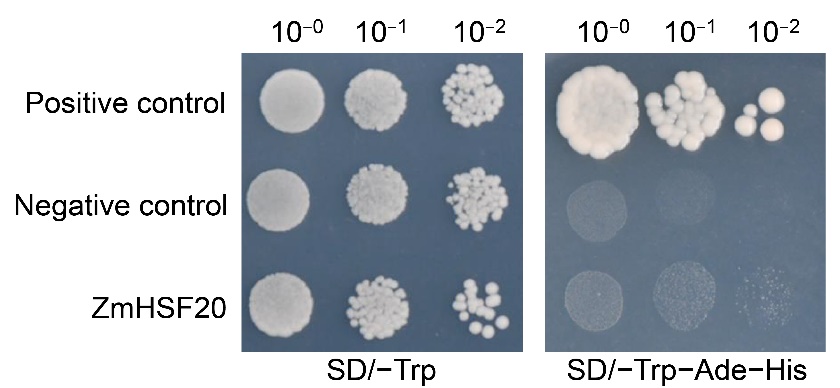

**Supplemental Figure. S3.** Transcriptional activation activity assay of ZmHSF20 in yeast. (Supports Figure 2) Positive control: pGBKT7-53 + pGADT7-T; Negative control: pGBKT7; ZmHSF20: pGBKT7-ZmHSF20.

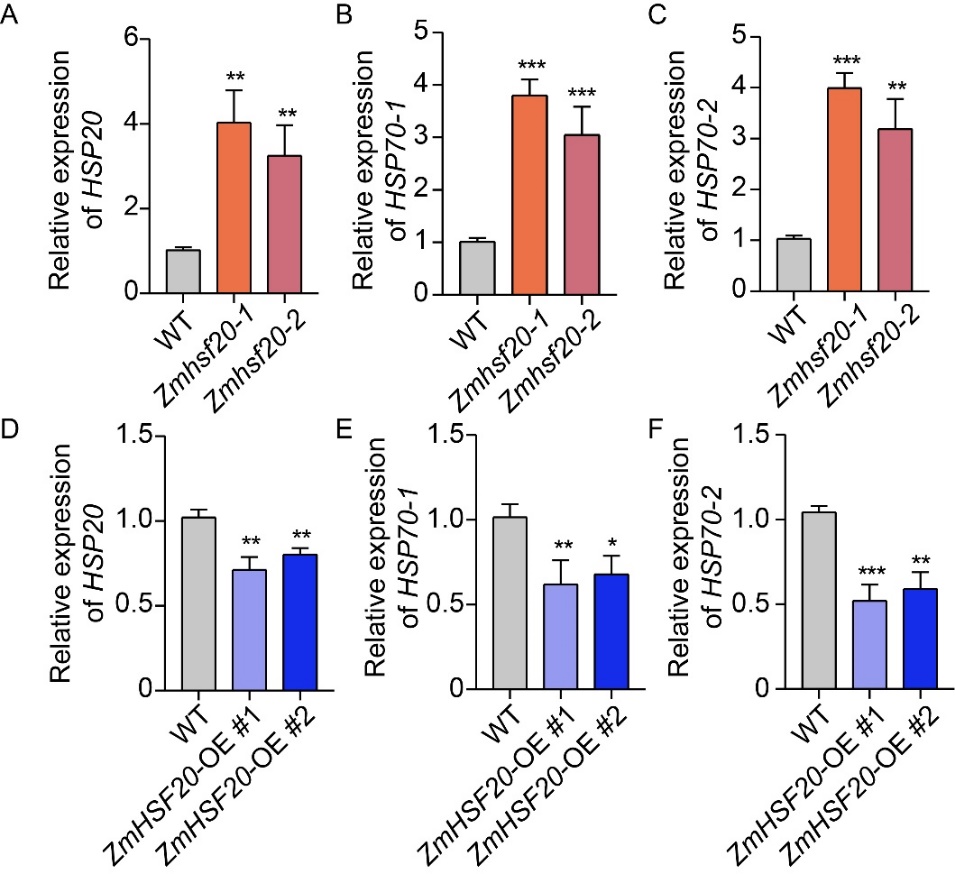

**Supplemental Figure. S4.** Effect of ZmHSF20 on the expression of *ZmHSP* genes. (Supports Figure 2) (A-F) Relative transcript levels of *ZmHSP20*, *ZmHSP70-1*, and *ZmHSP70-2* from V2 stage seedlings grown at 28℃/22℃, exposed to 45℃ for 1 day of the *Zmhsf20* mutant, *ZmHSF20*-OE, and WT. *ACTIN* was used as the internal control. The error bars are based on three independent experiments. The values are means ± SD (n = 3 independent experiments). **P* < 0.05, ***P* < 0.01, ****P* < 0.001, one-way ANOVA.

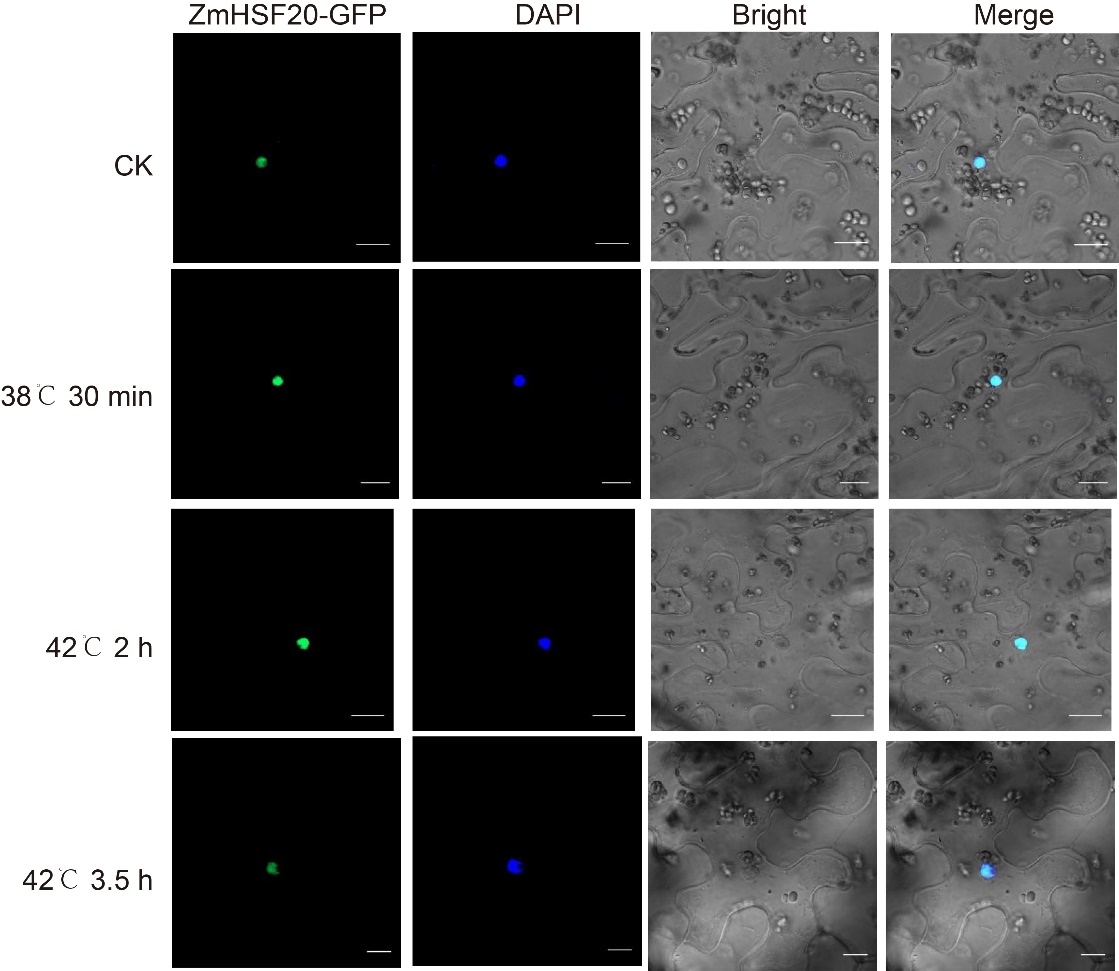

**Supplemental Figure. S5.** Subcellular localization of ZmHSF20. (Supports Figure 2) The indicated constructs were infiltrated into the leaves of *Nicotiana* *benthamiana* plants and observed under normal growth conditions or heat stress conditions stained with 4’,6-Diamidino-2-phenylindole (DAPI). Scale bar, 20 μm.

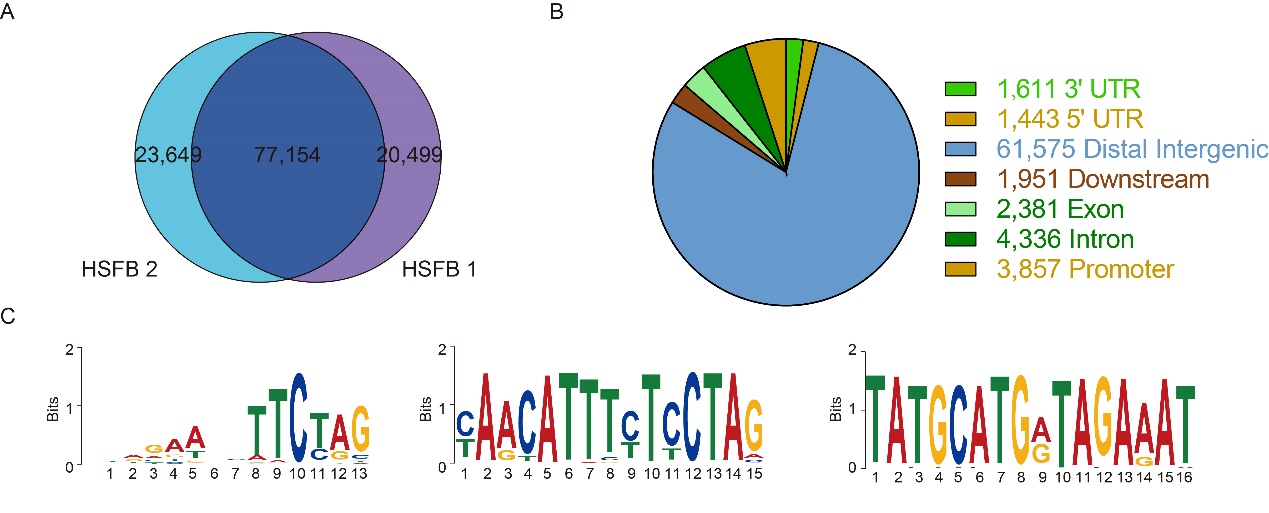

**Supplemental Figure. S6.** DAP-seq analysis. (Supports Figure 3) (A) Genome-wide identification of ZmHSF20 binding sites through DAP-seq. DAP-seq using two biological replicates reveals 77,154 high-confidence ZmHSF20 binding peaks. (B) Distribution of ZmHSF20 binding peaks across genomic features. (C) DNA logos of enriched DNA-binding sites for ZmHSF20 as determined by Homer. The core sequence of “nnGAAnnTTCnn” was substantially enriched among the ZmHsf20 binding regions and was named ZmHSE (mazie Heat Shock Element).

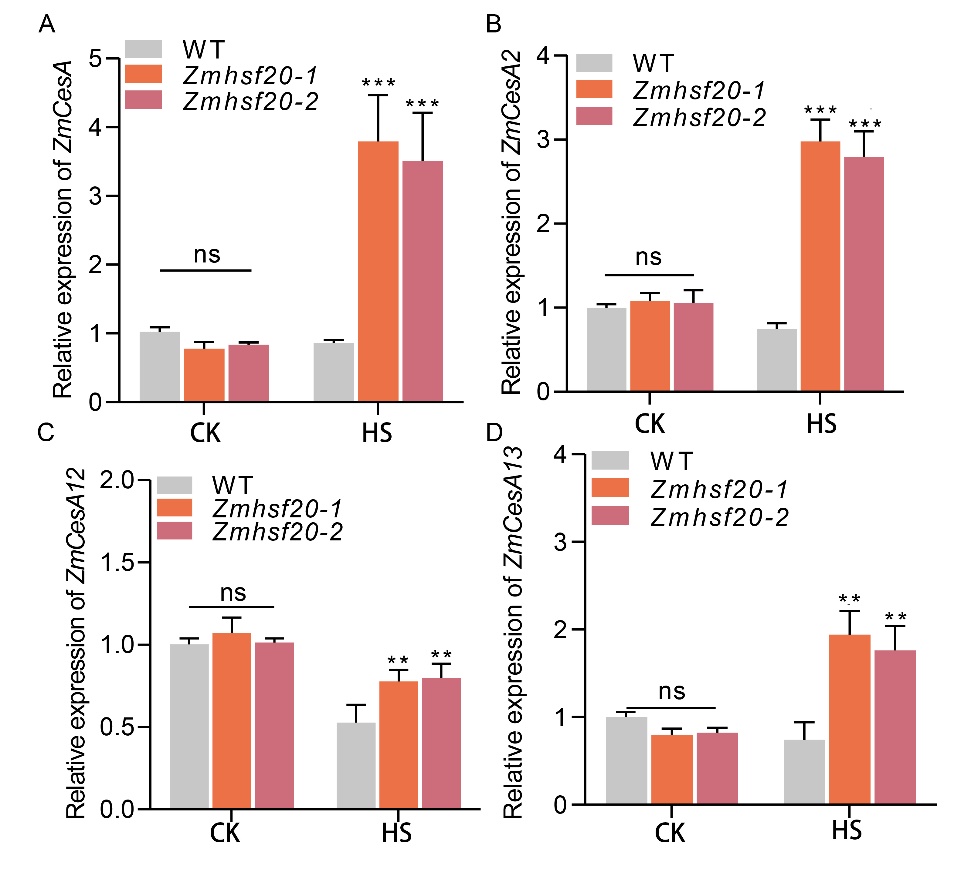

**Supplemental Figure. S7.** Effect of ZmHSF20 on the expression of *ZmCesA* genes. (Supports Figure 3) (A-D) Relative transcript levels of *ZmCesA*, *ZmCesA2*, *ZmCesA12*, and *ZmCesA13* in the leaves of V2 stage seedlings of *Zmhsf20* mutants and WT grown under normal conditions or after heat treatment for 24 h. *ACTIN* was used as the internal control. The error bars are based on three independent experiments. The values are means ± SD (n = 3). ***P* < 0.01, ****P* < 0.001, one-way ANOVA.

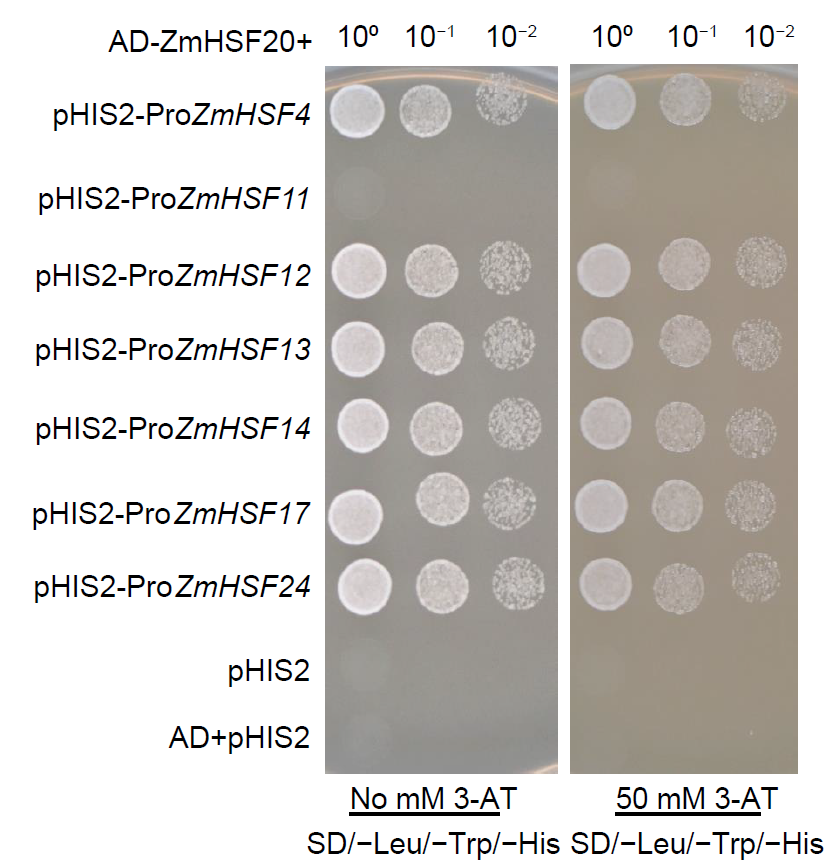

**Supplemental Figure. S8.** A yeast one-hybrid (Y1H) assay displaying direct binding of ZmHSF20 to the promoters of *ZmHSF4*, *ZmHSF13*, *ZmHSF17*, *ZmHSF24*, *ZmHSF14*, *ZmHSF12*, and *ZmHSF11*. (Supports Figure 4) Yeast clones were grown on synthetic defined (SD) medium lacking Leu, Trp, and His alone or containing 50 mM 3-amino-1,2,4-triazole (3-AT).

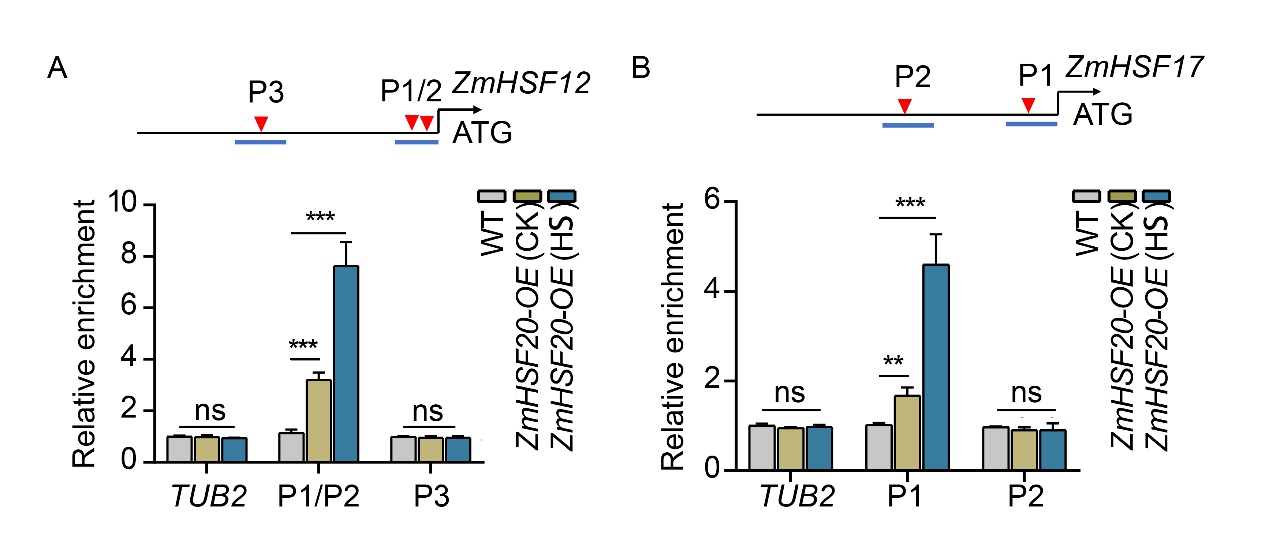

**Supplemental Figure. S9.** ZmHSF20 binds to the *ZmHSF12* and *ZmHSF17* promoters in vivo. (Supports Figure 4) (A) (B) CUT&Tag assay showing the binding of ZmHSF20 to the *ZmHSF12* and *ZmHSF17* promoters in vivo. WT and *ZmHSF20*-OE seedlings were exposed to heat treatment at 45℃ for 24 h or maintained at 28℃/22℃. CUT&Tag was performed using an anti-MYC antibody. Immunoprecipitated DNA was quantified by qPCR using primers specific to regions within the *ZmHSF12* or *ZmHSF17* locus. The relative enrichment of ZmHSF20 binding to the *ZmHSF12* and *ZmHSF17* promoters was normalized to *TUBULIN2* (*TUB2*). Each experiment was performed at least three times with similar results. The values are means ± SD (n = 3 independent experiments). ns, not significant. ***P* < 0.01, ****P* < 0.001, one-way ANOVA.

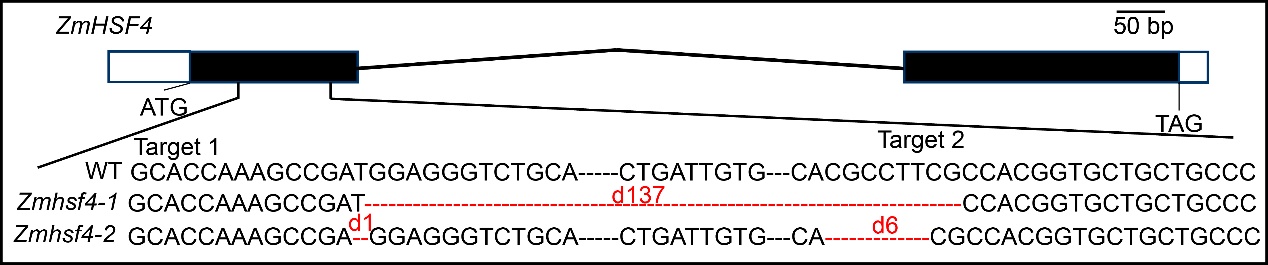

**Supplemental Figure. S10.** Construction of CRISPR/Cas9-based *Zmhsf4* knockout transgenic plants. (Supports Figure 4) Two sgRNAs that specifically target *ZmHSF4* were used, leading to the identification of two mutants, *Zmhsf4*-1 and *Zmhsf4*-2. black rectangles exons, white rectangles Un-Translated Regions (UTRs) and horizontal black lines rectangles introns.

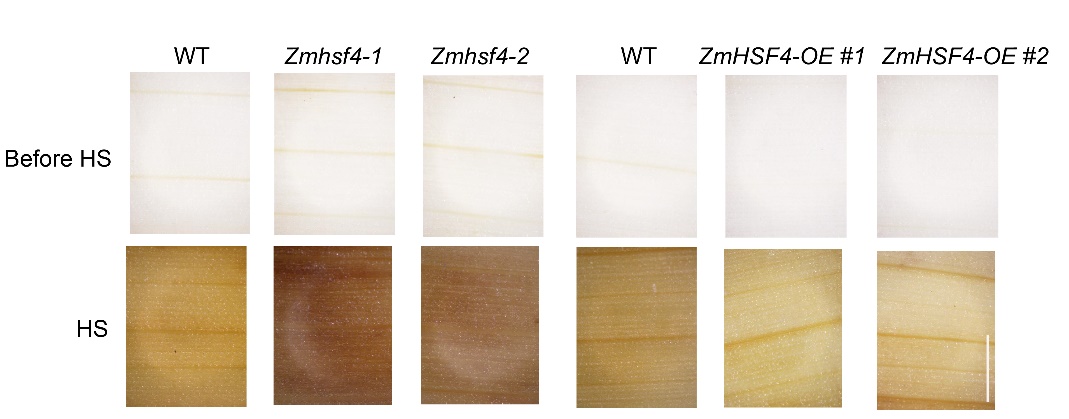

**Supplemental Figure. S11.** Heat tolerance is modulated by *ZmHSF4*. (Supports Figure 4) Representative photographs of the leaves of leaves from V2 stage seedlings grown at 28℃/22℃ exposed at 45℃ for 1 day and stained with DAB, scale bar = 1.5 mm.

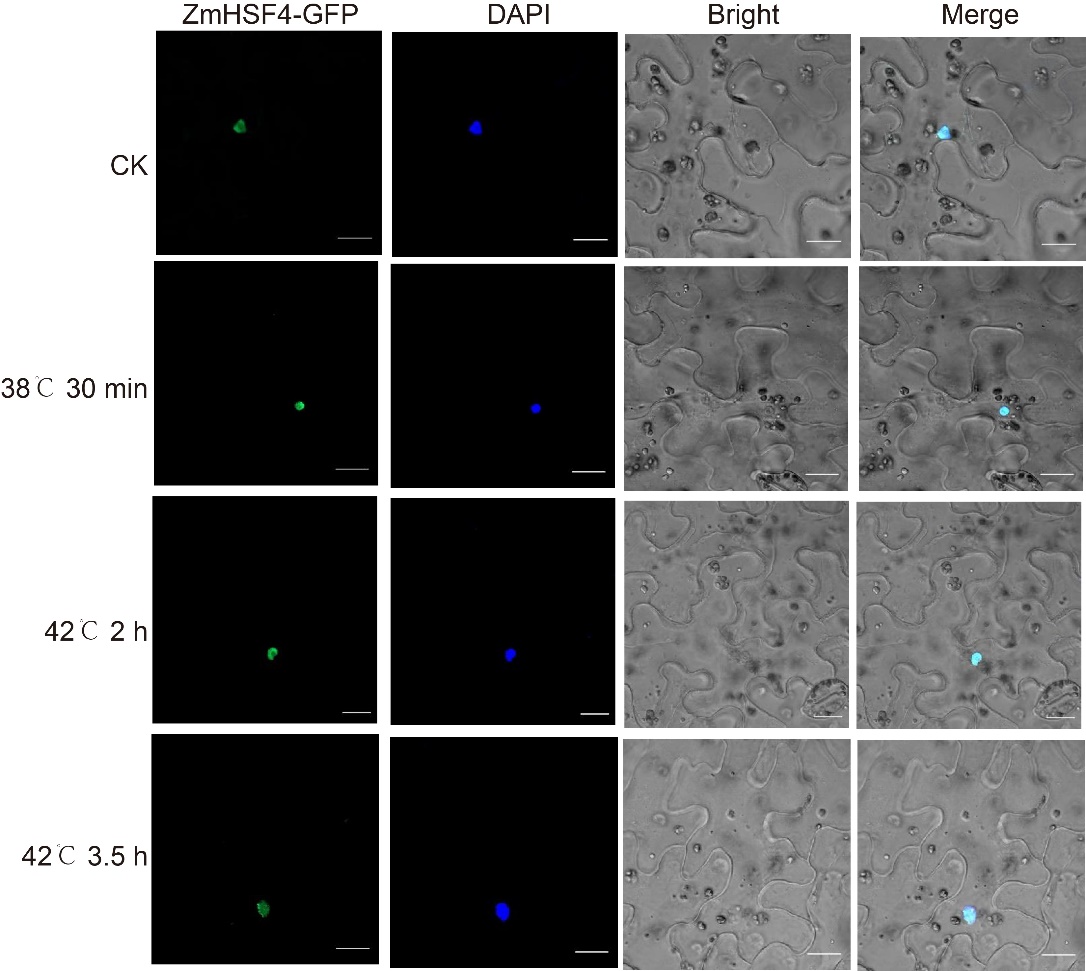

**Supplemental Figure. S12.** Subcellular localization of ZmHSF4. (Supports Figure 4) The indicated constructs were infiltrated into the leaves of *Nicotiana* *benthamiana* plants and observed under normal growth conditions or heat stress conditions stained with 4’,6-Diamidino-2-phenylindole (DAPI). Scale bar, 20 μm.

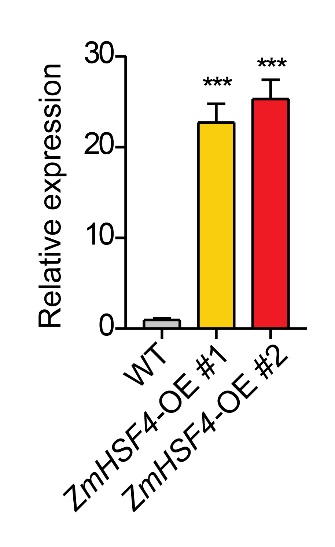

**Supplemental Figure. S13.** *ZmHSF4* expression level in overexpression lines. (Supports Figure 4) Relative transcript levels of *ZmHSF4* in the leaves of V2 stage seedlings of *ZmHSF4*-OE and WT. *ACTIN* was used as the internal control. The error bars are based on three independent experiments. The values are means ± SD (n = 3 independent experiments), ****P* < 0.001, one-way ANOVA.

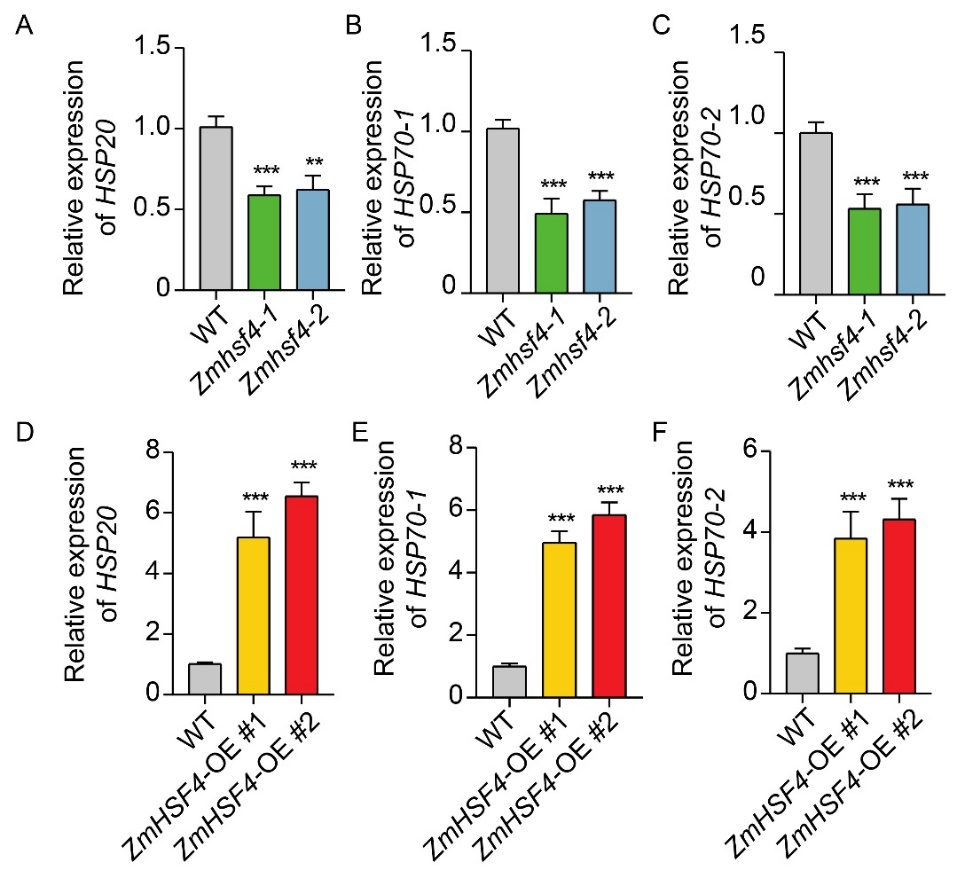

**Supplemental Figure. S14.** Effect of ZmHSF4 on the expression of *ZmHSP* genes. (Supports Figure 4) (A-F) Relative transcript levels of *ZmHSP20*, *ZmHSP70-1*, and *ZmHSP70-2* in the leaves of V2 stage seedlings of *Zmhsf4* mutants, *ZmHSF4*-OE, and WT after heat treatment for 24 h. *ACTIN* was used as the internal control. The error bars are based on three independent experiments. The values are means ± SD (n = 3 independent experiments). ***P* < 0.01, ****P* < 0.001, one-way ANOVA.

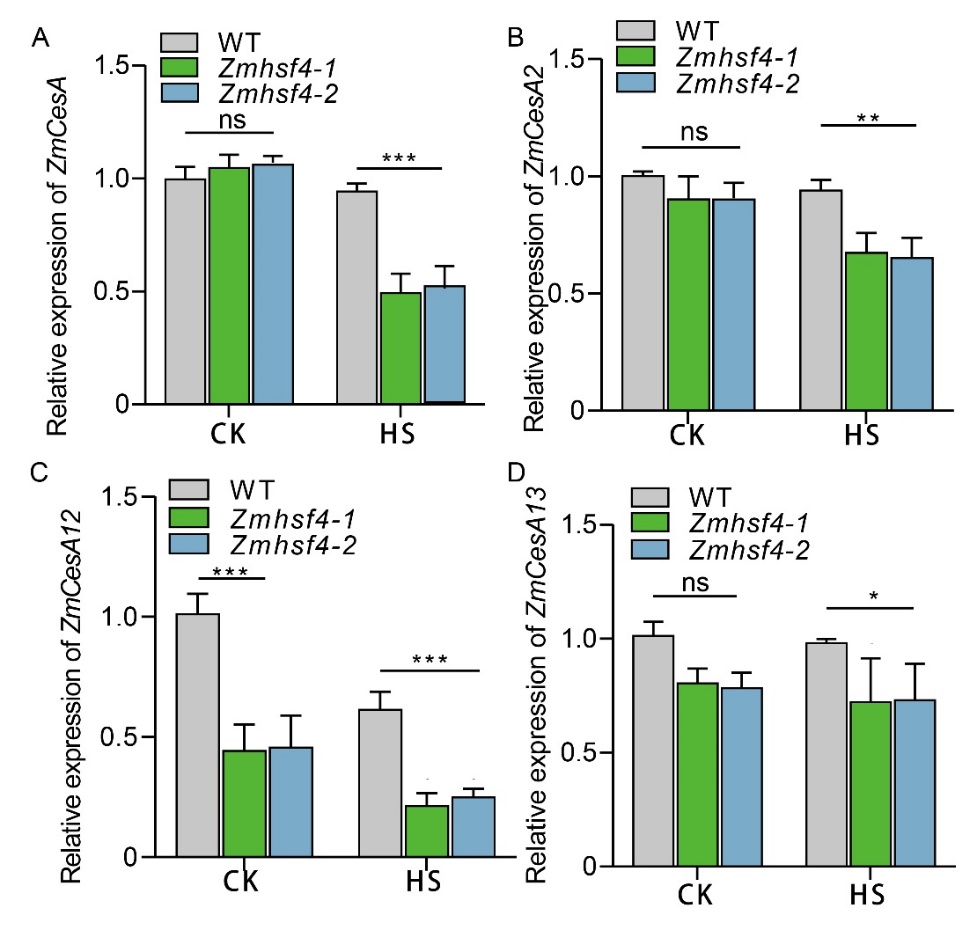

**Supplemental Figure. S15.** Effect of ZmHSF4 on the expression of *ZmCesA* genes. (Supports Figure 5) (A-D) Relative transcript levels of *ZmCesA*, *ZmCesA2*, *ZmCesA12*, and *ZmCesA13* in the leaves of V2 stage seedlings of *Zmhsf4* and WT grown under normal conditions or after heat treatment for 24 h. *ACTIN* was used as the internal control. The error bars are based on three independent experiments. The values are means ± SD (n = 3 independent experiments). **P* < 0.05, ***P* < 0.01, ****P* < 0.001, one-way ANOVA.

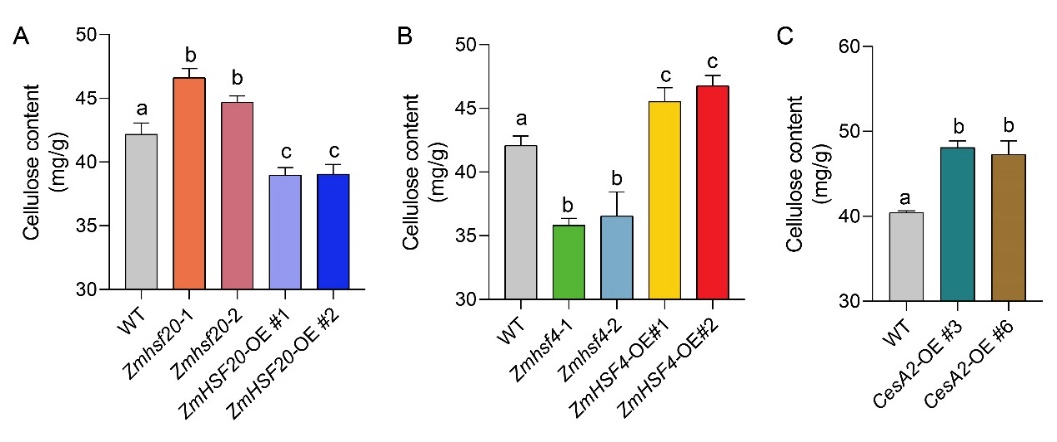

**Supplemental Figure. S16.** Measurement of cellulose contents. (Supports Figure 5) (A-C) Cellulose contents in WT, *ZmHSF20-OE*, *ZmHSF4-OE*, and *ZmCesA2-OE* lines under 45℃ heat stress for 1 day. Different lowercase letters indicate statistically significant differences (adjusted P < 0.05, one-way ANOVA).
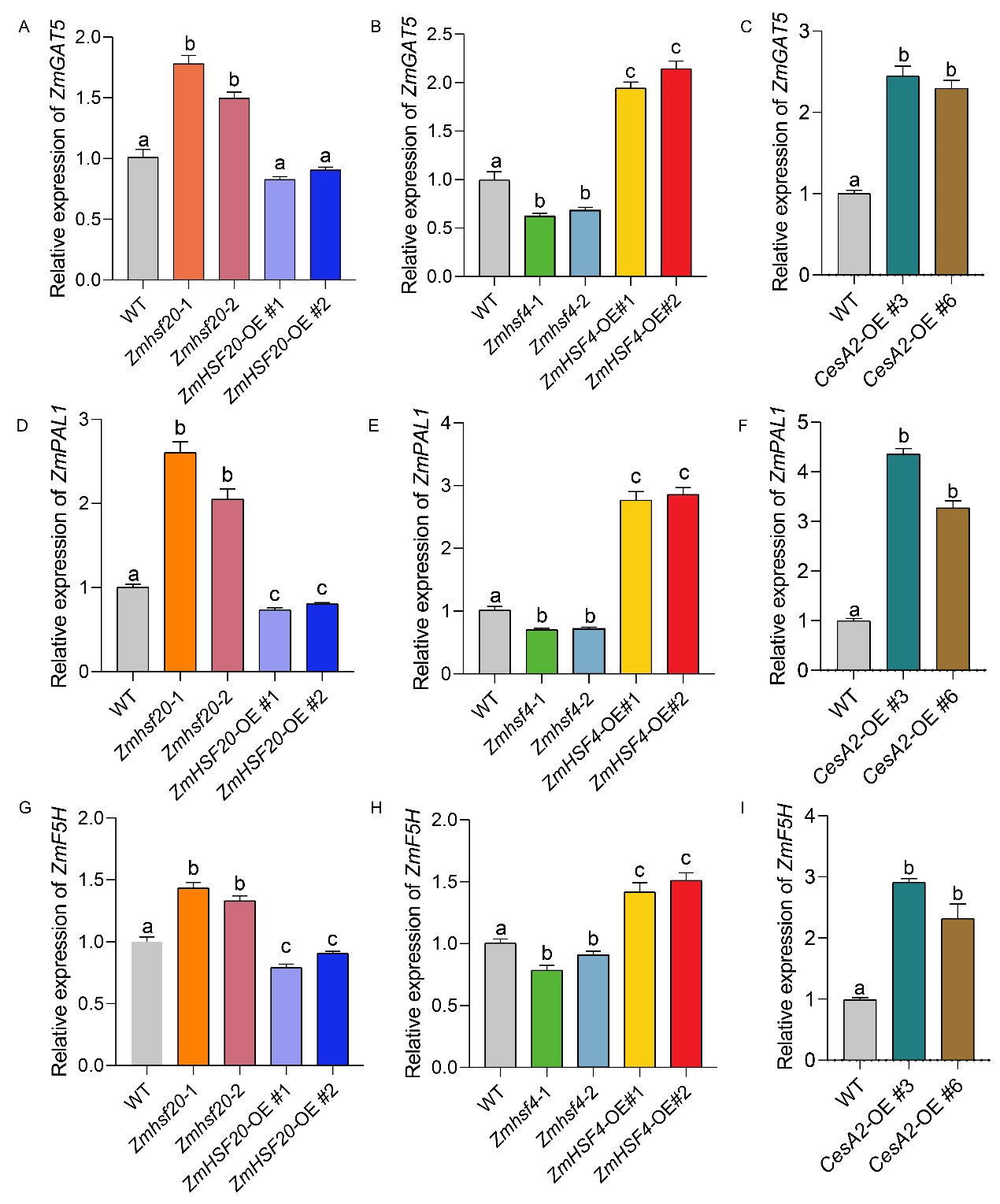

**Supplemental Figure. S17.** Effect of ZmHSF20, ZmHSF4 and ZmCesA2 on the expression of cell-wall regulated genes. (Supports Figure 5) (A-I) Relative transcript levels of *ZmGAT5*, *ZmPAL1*, and *ZmF5H* in the leaves of V2 stage seedlings of *Zmhsf20*, *Zmhsf4* mutants, WT, and *ZmCesA2*-OE lines after heat treatment for 24 h. *ACTIN* was used as the internal control. The error bars are based on three independent experiments. The values are means ± SD (n = 3 independent experiments). Different lowercase letters indicate statistically significant differences (adjusted *P* < 0.05, one-way ANOVA).

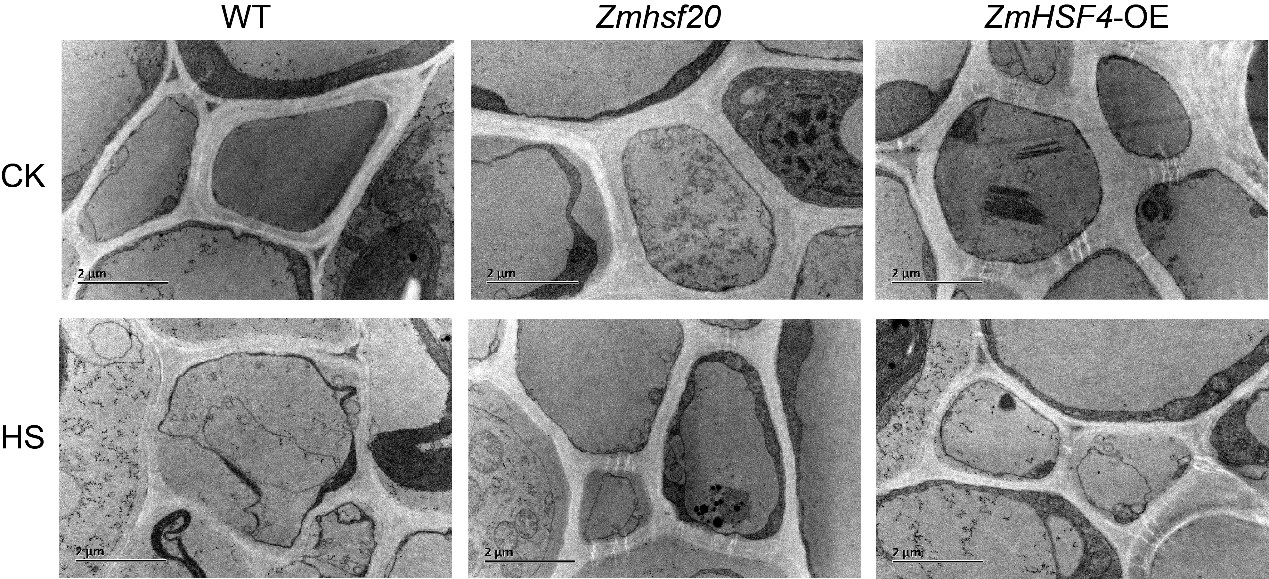
 **Supplemental Figure. S18.** Detection of the Structure of cell walls of WT, *Zmhsf20* mutants, and ZmHSF4-OE lines under normal and HS condition. (Supports Figure 5) The leaves from V2 stage seedlings grown at 28℃/22℃ (CK) or exposed to 45℃ for 2 days (HS) of WT, *Zmhsf20* mutants and *ZmHSF4-OE* via transmission electron microscope. Bars = 2 μm.

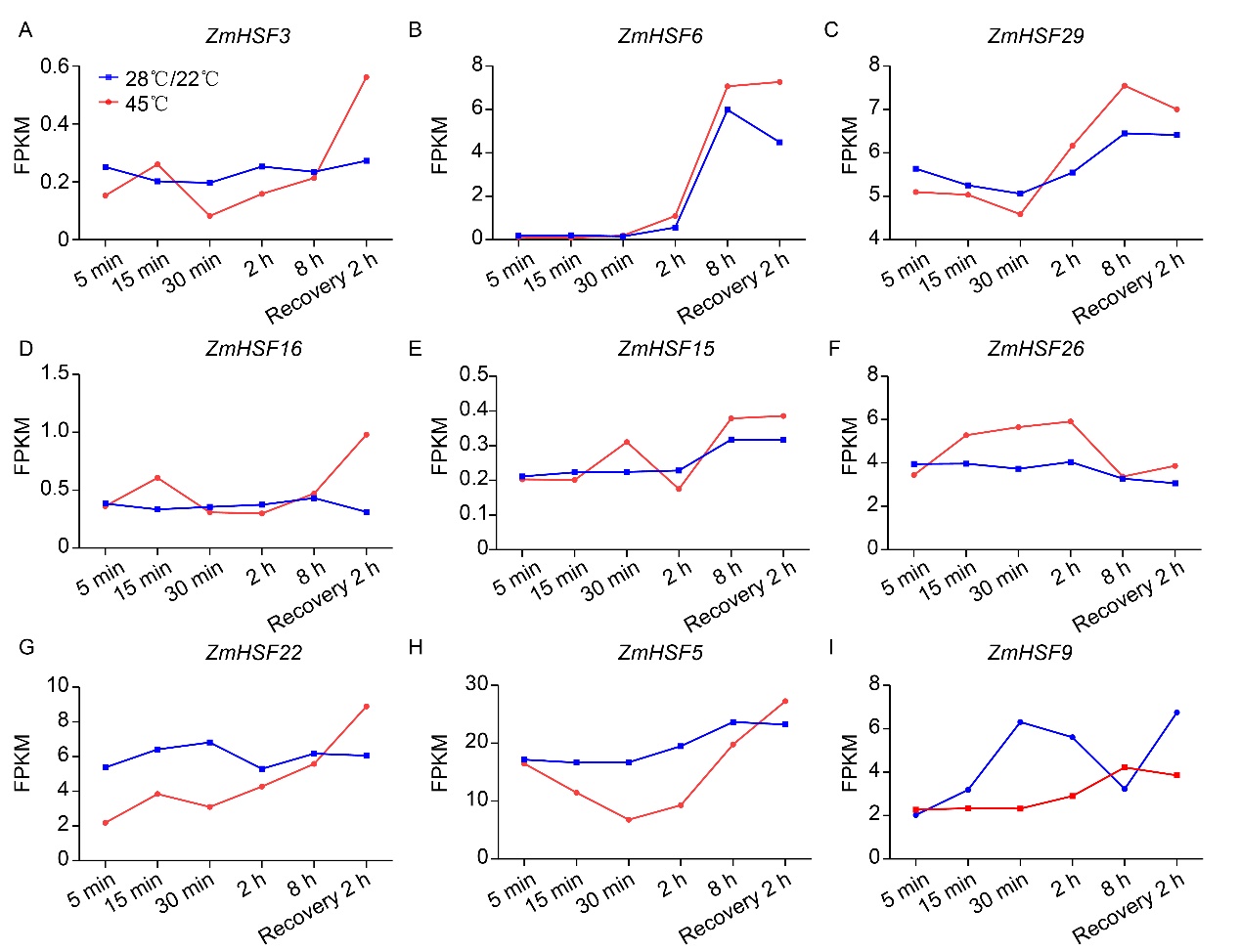

**Supplemental Figure. S19.** *ZmHSF* genes expression under 45℃ treatment no response or response weakly. (Supports Figure 1) (A-I) Expression pattern of *ZmHSF3* *ZmHSF6*, *ZmHSF29*, *ZmHSF16*, *ZmHSF15*, *ZmHSFf26*, *ZmHSF22*, *ZmHSF5* and *ZmHSF9* during heat treatment.

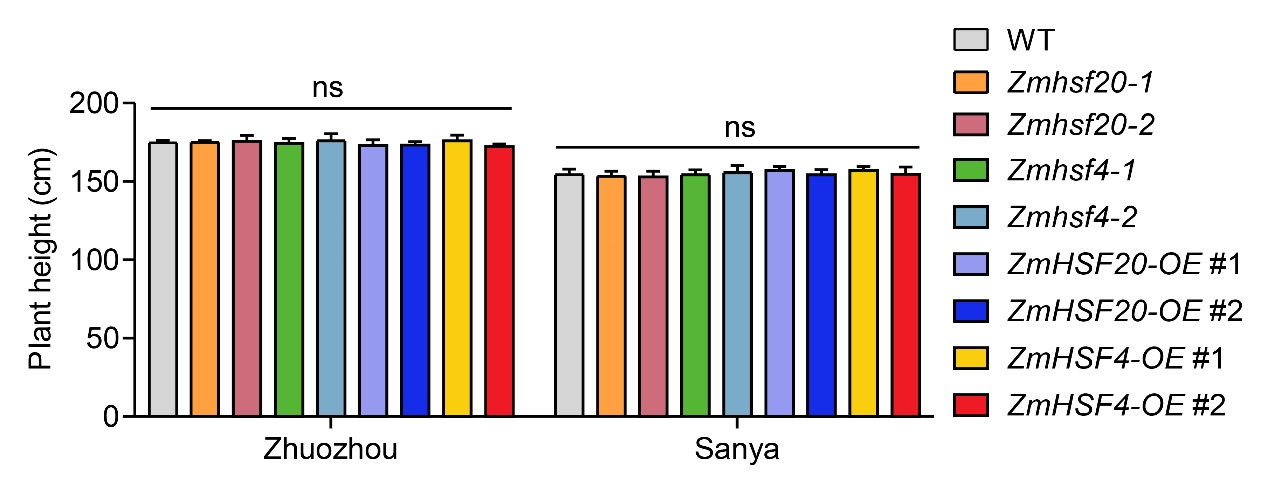

**Supplemental Figure. S20.** Plant height of different genotypes. (Supports Figure 5) Plant height of WT, *Zmhsf20*-1, *Zmhsf20*-2, *Zmhsf4*-1, *Zmhsf4*-2, *ZmHSF20*-OE #1, *ZmHSF20*-OE #2, *ZmHSF4*-OE #1, and *ZmHSF4*-OE #2 planted in two field stations in 2023. Blue bars, field data in Zhuozhou; green bars, field data in Sanya. Data represent means ± SD, ns, not significant.

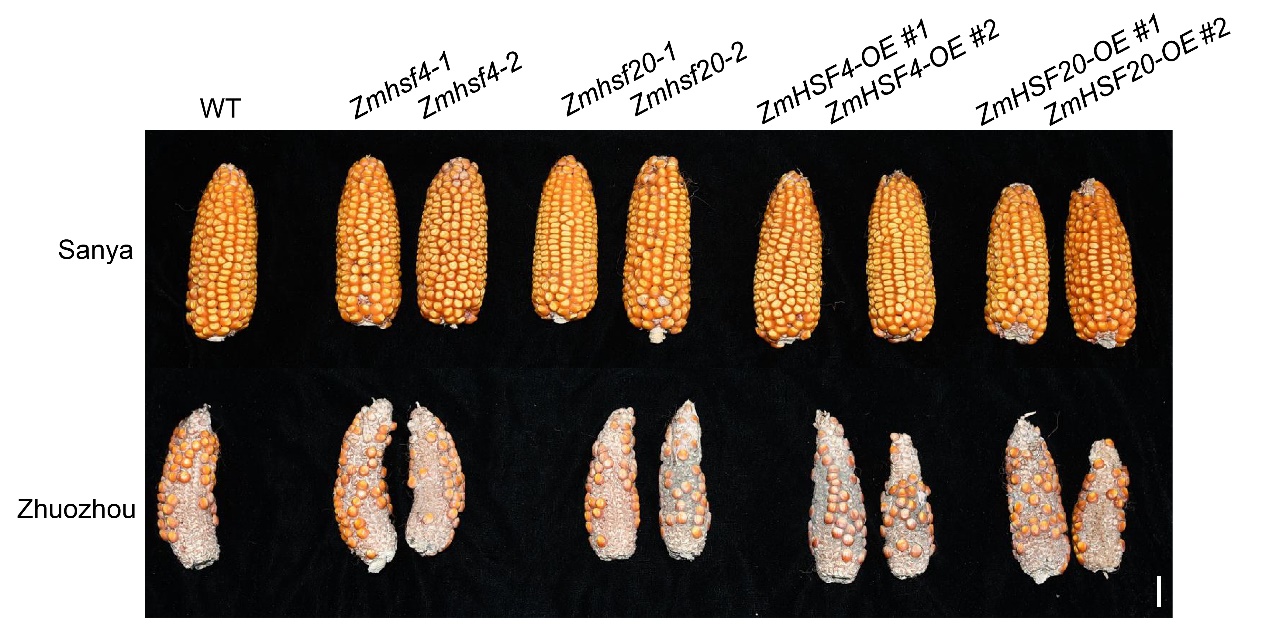

**Supplemental Figure. S21.** Representative photographs of ears of different maize genotypes planted in two field stations in 2023, (Supports Figure 5) Scale bar, 2 cm.
